# Transcriptional characterization of cocaine withdrawal versus extinction within nucleus accumbens

**DOI:** 10.1101/2024.03.12.584637

**Authors:** Freddyson J. Martínez-Rivera, Leanne M. Holt, Angélica Minier-Toribio, Molly Estill, Szu-Ying Yeh, Solange Tofani, Rita Futamura, Caleb J. Browne, Philipp Mews, Li Shen, Eric J. Nestler

**Author notes:** Co-correspondence Freddyson J. Martinez-Rivera, PhD Leanne M. Holt, PhD Eric J. Nestler, MD, PhD. Department of Neuroscience, McKnight Brain Institute, College of Medicine, University of Florida, Gainesville, Florida, USA. Co-first authors (Freddyson J. Martínez-Rivera, Leanne M. Holt).

## Abstract

Substance use disorder is characterized by a maladaptive imbalance wherein drug seeking persists despite negative consequences or drug unavailability. This imbalance correlates with neurobiological alterations some of which are amplified during forced abstinence, thereby compromising the capacity of extinction-based approaches to prevent relapse. Cocaine use disorder (CUD) exemplifies this phenomenon in which neurobiological modifications hijack brain reward regions such as the nucleus accumbens (NAc) to manifest craving and withdrawal-like symptoms. While increasing evidence links transcriptional changes in the NAc to specific phases of addiction, genome-wide changes in gene expression during withdrawal vs. extinction (WD/Ext) have not been examined in a context- and NAc-subregion-specific manner. Here, we used cocaine self-administration (SA) in rats combined with RNA-sequencing (RNA-seq) of NAc subregions (core and shell) to transcriptionally profile the impact of experiencing withdrawal in the home cage or in the previous drug context or experiencing extinction training. As expected, home-cage withdrawal maintained drug seeking in the previous drug context, whereas extinction training reduced it. By contrast, withdrawal involving repetitive exposure to the previous drug context increased drug-seeking behavior. Bioinformatic analyses of RNA-seq data revealed gene expression patterns, networks, motifs, and biological functions specific to these behavioral conditions and NAc subregions. Comparing transcriptomic analysis of the NAc of patients with CUD highlighted conserved gene signatures, especially with rats that were repetitively exposed to the previous drug context. Collectively, these behavioral and transcriptional correlates of several withdrawal-extinction settings reveal fundamental and translational information about potential molecular mechanisms to attenuate drug-associated memories.

## Introduction

Substance use disorder (SUD) is characterized by neurobehavioral adaptations that underlie devastating health and social problems ^1^. These adaptations are thought to be the driving force for relapse in humans with SUD. Remission often fails because withdrawal-associated symptoms become overwhelming and increase the propensity for relapse ^2–5^. While extinction-based therapies and adjunct medication have been used to manage withdrawal symptoms ^6–9^, these approaches show limited success, perhaps due to the lack of understanding of the fundamental molecular mechanisms involved.

Central to SUD-associated adaptations, the nucleus accumbens (NAc) and its two subregions (core and shell) signal specific information and exhibit orchestrated transcriptional changes in a time-, phase-, and context-dependent manner ^10–14^. It is suggested that the NAc core drives drug-seeking and -taking behaviors, whereas the NAc shell acts as an integrative hub for saliency and extinction processes ^15–17^. However, to date, the transcriptional changes in the NAc core and shell during withdrawal and extinction (WD/Ext) have not yet been characterized in drug self-administration (SA) models.

Here, we used two different withdrawal modalities after chronic cocaine SA in rats ^18^, whereby forced abstinence was achieved with repetitive context exposure (in the absence of drug, levers, or cue) or with home-cage confinement, and compared results during withdrawal to those obtained after full extinction conditions (context/levers/cue). This approach was combined with genome-wide RNA-sequencing (RNA-seq) of the NAc core and shell separately. Bioinformatic analyses were used to reveal gene expression patterns, biological functions, potential upstream regulators, and co-expression modules of gene networks in an NAc subregion-specific manner across behavioral conditions. To clinically leverage our transcriptomic findings, we integrated the present results with an available transcriptomic dataset of NAc samples from patients with cocaine use disorder (CUD) ^19^. Understanding the behavioral and transcriptional correlates of different WD/Ext settings will provide actionable information to better manage relapse triggers and achieve long-term abstinence.

## Materials and Methods

### Animals

Adult male Long-Evans rats (8-12 weeks; Charles River Laboratories) were paired-housed on a 12 h reverse light-dark cycle (lights on at 19:00) with food and water ad libitum. Drug SA experiments were executed during the dark phase (∼13:00-18:30) under red light illumination. The day before SA started (see below), rats were restricted to 18 g/day of standard laboratory chow for the rest of the study. All procedures were approved by the Institutional Animal Care and Use Committee (IACUC) of the Icahn School of Medicine at Mount Sinai, and the Association for Assessment and Accreditation of Laboratory Animal Care (AAALAC).

### Jugular vein catheterization (JVC)

Rats were anesthetized with isoflurane inhalant gas (5%; Patterson Vet., CO) in an induction chamber and positioned on a surgical pad with a face mask delivering 2-3% isoflurane for anesthesia maintenance. Rats were then prepared for the implantation of chronic indwelling catheters (in-house made) ^20,21^. Catheter tubing (0.012“ ID x 0.025” OD; Fisher Sci., NH) connected to a cannula (C313G-5UP; Plastic 1, VA / Protech International, TX) with a mesh (McMaster-Carr, IL) base secured with dental cement (Patterson Vet., CO) and inserted (2.5 cm) into the right jugular vein and held in place with silk threads (Fisher Sci.). This was followed by skin sutures (Patterson Vet., CO) and adhesive glue (Fisher Sci., NH). After surgery, rats were housed individually and their catheters were flushed with 0.1 ml heparinized saline (30 U) containing ampicillin (5 mg/ml; Patterson Vet., CO) once a day during the recovery phase (4-6 days). Carprofen (5 mg/kg Patterson Vet., CO) was subcutaneously administered as an analgesic. The day before SA started (see below), catheters were tested for patency using methohexital (5 mg/kg; Cardinal Health, OH) followed by 0.1 ml of heparinized saline (30 U) containing ampicillin. This procedure was repeated as needed.

### Drug

Cocaine HCl (NIDA) was dissolved in saline during the SA training (acquisition phase).

### Operant boxes

Cocaine was delivered from a syringe pump (Med Associates, VT) with a 10 ml syringe connected to a swivel via polyethylene-50 tubing (Plastic 1, VA) on both ends. At the exit end of the swivel, the tubing was protected with a metal mesh and connected to the cannula (catheter) on the back of the rats. The operant boxes were located inside sound-attenuating cabinets and controlled by an interface system (Med Associates, VT) ^20,21^.

### SA phases (acquisition, Test 1, withdrawal, extinction, and Test 2)

*Acquisition:* Rats were trained to self-administer cocaine (0.8 mg/kg/infusion) on a FR1 ratio during 3 h sessions over 10 days ^20–22^. Infusions (rewards) were signaled by a cue light above the active (correct) lever followed by a 20 s timeout period in which the cue light remained illuminated and levers extended. During the timeout, levers remained extended but without consequence. Inactive (incorrect) lever presses were recorded but had no consequences. During this phase, the catheters were flushed before and after each session with heparinized saline (0.1 ml; 30 U).

*Test 1:* Two days after the last acquisition day, rats were placed in the operant boxes under full extinction conditions (exposed to context/levers/cue but no drug consequences) for 30 min. The next day, rats were separated into three different WD/Ext groups: home-cage, context withdrawal, or full extinction: 1) *Context exposure (Ctx):* Rats were exposed daily to the previous drug context (operant boxes for 1 h sessions) but without levers, cues, or drug over the course of 14 days. 2) *Home cage (HC):* Rats were kept in their home cages (no context, levers, cues, or drug) and handled daily in the behavioral rooms for 14 d (while groups 1 and 3 were in the operant boxes). 3) *Extinction training (Ext):* Rats were placed daily in the operant boxes for 1 h sessions with full access to the context, levers, and cue light but without drug infusions over 14 days.

*Test 2:* Two days after the last WD/Ext session, rats were placed in the operant boxes under full extinction conditions for 30 min and lever presses were recorded.

### Tissue collection

Within ∼30 min after Test 2, rats were decapitated, and the brains were extracted and placed on ice-cold PBS (∼2 min) and sectioned into 1 mm thick coronal sections using a prechilled brain matrix. Bilateral brain punches of the NAc core and shell (15 and 18 gauge, Scientific Commodities, AZ) were collected (coordinates AP; +2.76 to +0.70) ^23^, placed in Eppendorf tubes, and flash frozen on dry ice and stored on -80°C until further analyses.

### RNA isolation and library preparation

Total RNA was purified from bilateral NAc core and shell separately using the Direct-zol RNA kit following the manufacturer’s instructions (Zymo Research, CA), including DNase to remove genomic DNA. RNA integrity number (RIN: 6.5-9) and concentration (5-50 ng/µl; 260/280 values >1.7) were evaluated using the Agilent 2100 Bioanalyzer (Agilent Technologies, CA) and Nanodrop (Thermo-fisher, MA). For library preparations, we used SMARTer Stranded Total RNA-seq kit v3 - Pico Input Mammalian (Takara Bio, CA). All libraries were in-house tested for quality control (averaging 23 ng/µL, Qbid; range peak size 300-500 bp, Bioanalyzer) ^24,25^.

### Sequencing and differential expression gene analysis

Sequencing was performed by Genewiz/Azenta (MA) with library quality control performed on TapeStation (TS) and Qbid (range peak size 364-590 bp, TS; 18-92 ng/µL, Qbid). Sequencing was conducted on Illumina NovaSeq instruments with a 2×150 bp paired-end read configuration. All samples (84 in total) were multiplexed and run concurrently to produce 30M paired-end reads per sample. Raw sequencing reads were mapped to the rn6 rat genome (with annotation GCF_000001895.5) using the NGS-Data-Charmer pipeline (https://github.com/shenlab-sinai/NGS-Data-Charmer). In brief, reads were trimmed to remove adaptor and low-quality sequences and then aligned to the reference genome. Duplicate alignments were removed before gene counting and downstream analysis. Counts of mapped reads were obtained using feature Counts against the GENCODE vM25 annotation. After quality control was assessed and poor-quality samples removed, differential expression analysis was performed on genes with a minimum mean count of 6 (to remove low-count genes), which yielded a total list of 29,888 genes. Differential expression was performed in R version 4.0.2 using the DESeq2 package. Unless otherwise specified, significance for differentially expressed genes (DEGs) was set at a Fold Change (logFC) of ± 25% and p<0.05.

### Biotype and cell type

Using the Ensembl *biomaRt* R package, resulting DEGs (annotated and unannotated) were filtered and categorized into different biotypes including protein-coding, pseudogenes, predicted, miRNA, rRNA, lncRNA, scRNA, snoRNA, and snRNA. For cell-type enrichment, DEG lists were categorized using an “enriched” measure from single-cell RNA-seq database comparisons ^26^, which classifies subsets of genes as being enriched in astrocytes, endothelial cells, microglia, neurons, oligodendrocytes, or oligodendrocyte precursor cells in the brain.

### Volcano plots and union heatmaps

Volcano plots were generated in R (v4.0.2) using ggplot (v3.4.2). Venn diagrams were generated using nVenn (PMID: 29949954) (http://degradome.uniovi.es/cgi-bin/nVenn/nvenn.cgi). Union heatmaps of each subregion (core and shell) were generated by creating a reference list of all DEGs present in each condition and merging the Log_2_FC of each condition with this reference list. Union heatmaps were then generated using Morpheus (https://software.broadinstitute.org/morpheus).

### Rank-rank hypergeometric overlap (RRHO2)

Stratified, two-sided RRHO2 plots were generated using the RRHO2 R package with standard settings (github.com/RRHO2/RRHO2) ^27^.

### Gene ontology and upstream regulator analysis

Gene ontology for “Biological Processes 2021” and “Molecular Functions 2021” databases were performed in R using the *enrichR* package with our filtered DEG lists as input (25% fold change (logFC) and p<0.05) ^19,28^. Ingenuity Pathway Analysis (IPA; Qiagen) was used to predict upstream regulators associated with DEGs (25% fold change (logFC) and p<0.05) for each experimental condition. Predicted regulators were exclusively considered as upstream regulators using the following and additional criteria: Benjamini-Hochberg-corrected p-value of p<0.05, where the regulators had to have a non-zero or non-null activation z-score ^29^. Consequent union heatmaps were generated entering the resulting regulators across conditions using Morpheus (https://software.broadinstitute.org/morpheus).

### Alluvial plots

Alluvial plots were generated to visualize the dynamic displacement of gene clusters across conditions and NAc subregions as previously used ^29^. Genes were categorized/clustered according to significance values (p<0.05) and mapped using the ggalluvial package in R, version 0.12.3.

### Motif enrichment analysis

We used the web-based toolkit Multiple Expectation Maximizations for Motif Elicitation (MEME) Suite for Simple Enrichment Analysis (SEA) for our experimental groups and NAc subregions. SEA identifies previously discovered motifs that are enriched in a set of input sequences (Bailey and Grant, 2021). As a result, multiple RNA or DNA binding proteins (including transcription factors) are provided based on their statistical significance including p-value, q-value (false discovery rate) ^30,31^, positional distribution, matches per sequences, motif logos, and other measures. Union heatmaps (Morpheus; https://software.broadinstitute.org/morpheus) were generated to highlight the top 30 motifs for each behavioral condition and merged by subregions to visualize the relationship of the resulting transcription factors across our variables.

### Gene co-expression network analysis

We used multiscale embedded gene co-expression network analysis (MEGENA) ^32–34^ to construct gene modules (networks) for each group of animals and NAc subregion. This approach reveals hub genes for each module (sunburst plots). In addition, an algorithm for the reconstruction of accurate cellular networks (ARACNE) ^35^ was used to identify likely transcriptional regulators. These analyses were generated using. ARACNe-AP, the “MEGENA” R package and the aesthetics were modified using the R package github.com/mw201608/sunburst.shiny ^32^.

### Comparisons with human RNA-seq CUD dataset

We leveraged recently published RNA-seq data of the NAc from human subjects who died from CUD ^19^. RRHO2 and heatmaps were generated to compare gene expression patterns between WD/Ext groups in rats and human CUD.

### Statistics

Behavioral data were plotted as mean ± standard error (SEM) and statistical significance was established as p< 0.05 using the Fisher exact test, t-tests, repeated measures, or mixed model (two-way) ANOVAs, followed by post hoc Sidak analysis (Prism 9). Percent change was described as ([active presses in Test 2 - active presses in Test 1] / total active presses) x 100.

Morpheus (https://software.broadinstitute.org/morpheus) was used to make heatmaps and perform hierarchical clustering analysis with default settings (one minus Pearson correlation and average linkage method). Gene expression data are plotted based on log2 FC and p-value thresholds as indicated throughout. Plots and drawings using Prism 9, biorender.com, the “tidyverse” R package (version 1.3.1), and github.com/eackermann/ratpack.

## Results

### Repeated context exposure in the absence of levers/cues/drug increases drug seeking

To investigate the transcriptional landscape of different forced abstinent scenarios, food-restricted and catheterized rats first underwent cocaine or saline SA followed by three different WD/Ext modalities: context or home-cage withdrawal or extinction training (Fig. 1A). Rats successfully acquired cocaine SA in daily 3-h sessions under an FR1 schedule (0.8 mg/kg/infusion) for 10 days with discrimination observed between the active and inactive levers (Figs. 1B; cocaine group: active vs. inactive presses; F_(1,92)_=298.7, p<0.001). We additionally confirmed successful acquisition by comparing the number of active lever presses between cocaine and saline groups (Fig. S1A and B) (Group; F_(1,47)_=94.45, p<0.001). Two days after the last acquisition session, rats exhibited drug-seeking behavior in a drug-free test (Test 1; t1) (Fig 1C; cocaine group: active vs. inactive presses; t_(92)_=10.98, p<0.001). Cocaine rats (Fig. 1C) additionally exhibited elevated seeking behavior compared to their saline counterparts (Fig. S1C; active presses; t_(75)_=11.86; p<0.001).

**Figure 1.**
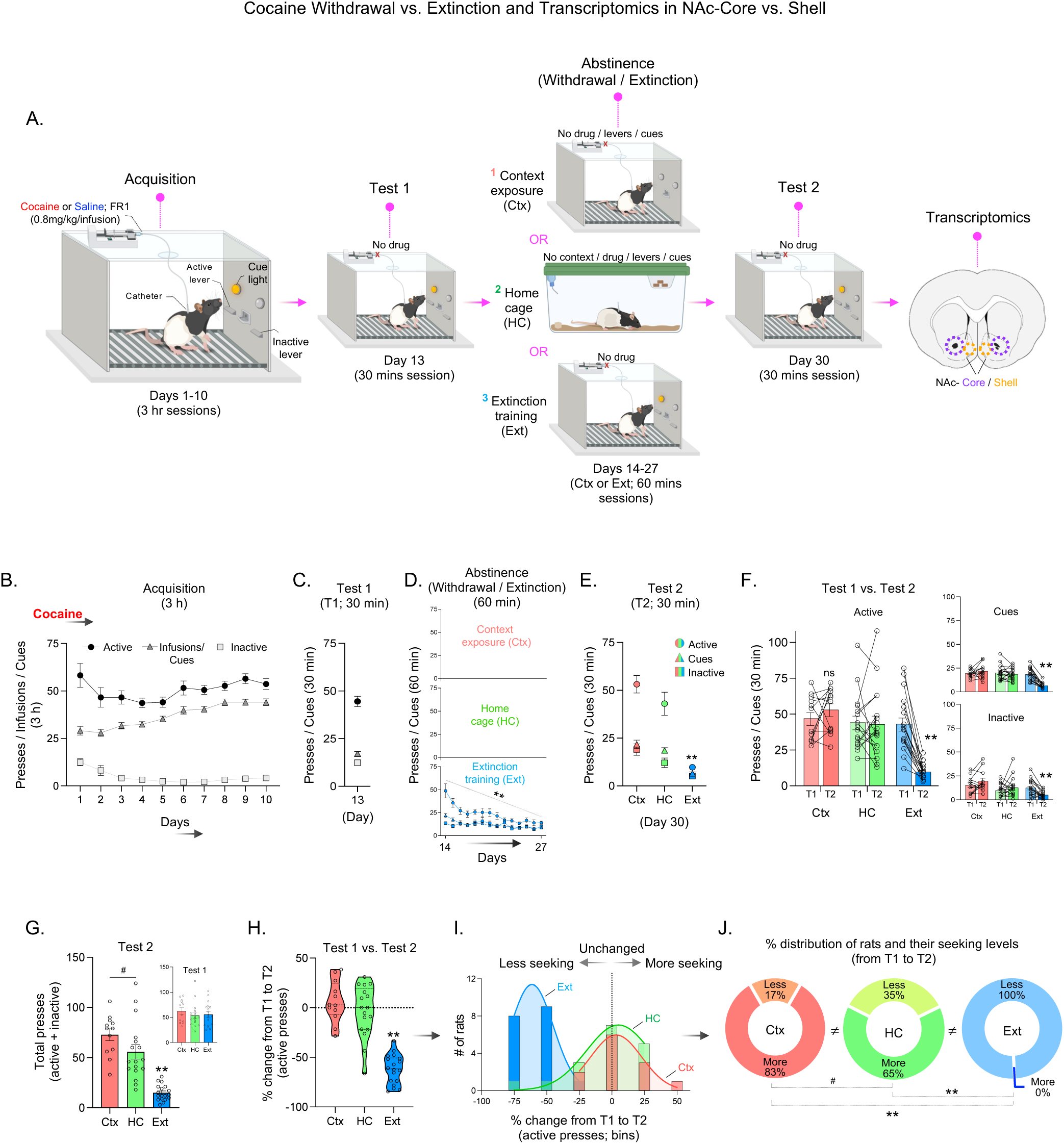
Context-exposed withdrawal increases cocaine-associated seeking. **A.** Schematic depicting experimental design; cocaine withdrawal/extinction (WD/Ext) modalities followed by transcriptomics (RNA-seq) in NAc subregions. **B-C.** Acquisition of cocaine SA and Test 1 (T1; drug-free). **D-E.** Separation of the experimental groups undergoing WD/Ext in the previous drug context (Ctx), in the home-cage (HC), or under full extinction conditions (Ext), followed by Test 2 (T2; drug-free). **F-G.** Comparison between Tests 1 and 2 displayed increased, unchanged, and reduced seeking behaviors in the Ctx, HC, and Ext groups, respectively. These effects were observed in active and inactive lever pressing (total lever pressing). **H-J.** Percent change analysis from Tests 1 to 2 confirms that Ctx-exposed rats display incubation-like behavior, while the HC and Ext groups remained stable or reduced their seeking, respectively. This interpretation was supported by the distribution of rats within the experimental groups. Cocaine, Ctx: n=12; HC: n=17; Ext: n=18. All data are shown as mean ± SEM. **p≤0.01, *p≤0.05, ^#^p≤0.10.

To probe adaptations after different WD/Ext paradigms, rats were separated into three subgroups: 1) placement back into the previous drug context (Ctx; context withdrawal), 2) home-cage (HC; home-cage withdrawal), or 3) under extinction conditions (Ext) (Fig. 1A and D). These groups were matched based on their performance during the acquisition and Test 1 phases (Fig. S1G; acquisition: active, F_(2, 44)_=0.22, p>0.05; inactive, F_(2, 44)_=3.92, p<0.05, Sidak post-hoc, day 1 (only; inset inactive) HC vs. Ext p<0.05; rewards; F_(2, 44)_=0.40, p>0.05; Test 1: F_(2, 132)_=0.63, p>0.05). In parallel, saline groups were concurrently separated into each experimental condition (Fig. S1D-F); these subgroups were also identical during acquisition and Test 1 (Figs. S1H; acquisition: active, F_(2, 27)_=0.36, p>0.05; inactive, F_(2, 27)_=0.87, p>0.05; rewards; F_(2, 27)_=0.83, p>0.05; Test 1: F_(2, 81)_=0.14, p>0.05). As expected, rats receiving full extinction training, including levers and cues, in the original drug context (Ext) reduced their seeking behavior after daily 1 h sessions over 14 days (Fig. 1D; active lever pressing over time: F_(5, 183)_=9.54, p<0.01; Sidak post-hoc, first vs. last day p=0.02). In contrast, no extinction occurred in saline conditions (Fig. S1D; time, F(6, 195)=2.46, p=0.03; Sidak post-hoc, all p>0.05). Rats from the Ctx groups (cocaine or saline) were exposed to the previous cocaine/saline context without any drug-associated stimuli, including any levers or cue lights. Rats from the HC group were also brought to the behavioral rooms daily to account for handling.

Two days after the last WD/Ext session, all three groups received another drug-seeking test (drug-free; Test 2). As anticipated, both the HC and Ctx groups showed elevated seeking behavior when compared with the Ext group (Fig. 1E; F_(2, 134)_=54.38, p<0.01, Sidak post-hoc, p<0.05), with no significant differences observed between the HC and Ctx groups (Fig. 1E, p>0.05; Sidak post-hoc). Some of these results were observed to a lesser extent in the saline counterparts for Ctx and HC groups (Fig. S1E; F_(2, 81)_=12.44, p<0.01, Sidak post-hoc, p<0.05). Nonetheless, such effects were dramatically higher in cocaine-exposed groups (as compared to saline counterparts) in both Tests 1 and 2 (Fig. S1I; Test 1: F_(1, 71)_=115.10, p<0.01; Test 2: F_(1, 71)_=57.85, p<0.01, Sidak post-hoc, p<0.01). Furthermore, when the Ext group (9.77 ± 1.23) is directly compared against its corresponding saline group (3.54 ± 0.87) in Test 2, a higher but not significant seeking behavior was still observed (Fig. S1I; Sidak post-hoc, p>0.05), highlighting the impact of previous cocaine experience and extinction training. These results were anticipated, given the exploratory nature of rats, and collectively support the importance of proper experimental controls for all behavioral conditions.

To further uncover potential and specific behavioral features within and between these WD/Ext modalities, we compared Tests 1 and 2 in cocaine or saline-exposed groups. Neither Ctx nor HC groups demonstrated statistically significant changes in seeking behaviors (active/inactive pressing and cues; Fig. 1F and insets) from Test 1 to 2 (Fig. 1F; active pressing: group, F_(2, 44)_=10.88, p<0.01; time, F_(1, 44)_=9.47, p<0.01, Sidak post-hoc, p>0.05). However, we observed increased seeking in Ctx rats (Fig. 1F, active lever presses; T1: 47.0 ± 4.5; T2: 53.2 ± 4.5, p>0.05), perhaps indicating facilitation in the development of incubation processes. Consistent with this interpretation, Ctx rats showed more total presses (active + inactive) during Test 2 compared to HC or Ext groups (Fig. 1G; F_(2, 44)_=30.91, p<0.01, Sidak post-hoc, Ctx vs. HC: p=0.08, Ctx vs. Ext: p<0.01), an effect that was not observed in Test 1 (Fig. 1G, inset; F_(2, 44)_=0.50, p>0.05) or directly replicated in saline conditions (Fig. S1J; Test 1: F_(2, 27)_=0.03, p>0.05; Test 2: F_(2, 27)_=5.3, p<0.05, Sidak post-hoc, Ctx vs. HC: p=0.85, Ctx vs. Ext: p<0.05, HC vs. Ext: p=0.06). As expected, the Ext group reduced their seeking (active/inactive pressing and cues; Fig. 1F and insets) in the same drug-associated context from Test 1 to Test 2 (Fig. 1F; active lever pressing: time, F_(1, 44)_=9.47, p<0.01, Sidak post-hoc, p<0.01). These effects were not recapitulated in saline groups (Fig. S1F and insets), except the Ctx saline group. However, cocaine groups still show higher seeking behavior as compared with saline groups (see Fig. S1I, bottom panel) despite the presses associated with the cue light presentation after a long omission break under saline and drug conditions (see Figs. S1I-J, bottom panels). This again underscores the relevance of appropriate control groups.

Subsequent analysis demonstrated subtle but important differences in the distribution of seeking behavior for each cocaine group after the WD/Ext phase. Unsurprisingly, Ext rats demonstrated a significantly negative percent change from Test 1 to Test 2 when compared to Ctx and HC groups, with no apparent differences between Ctx and HC (Fig. 1H; F_(2, 44)_=49.18, p<0.01; Sidak post-hoc, Ctx/HC vs. Ext: p’s<0.01; Ctx vs. HC: p>0.05). However, frequency distribution analysis revealed a higher proportion of Ctx rats showing elevated seeking (Fig. 1I; from Test 1 to Test 2: >-10% change). Rats were further characterized into “more seeking” or “less seeking” from Test 1 to Test 2 (>-10% or ≤-10% change). The specific proportions of rats showing more or less seeking were then compared with Fisher’s exact test, revealing a significant difference between Ctx and HC rats (Fig. 1I-J; p<0.01). Together, these results suggest that repeated context exposure during withdrawal worsens withdrawal-associated responses rather than alleviating them by promoting context extinction.

### Repeated context exposure in the absence of levers/cues/drug robustly alters the NAc core and shell transcriptome

To transcriptionally link our behavioral findings of various withdrawal and extinction conditions to the brain’s reward circuitry, RNA-seq was performed on NAc core and shell dissections collected 30 min after Test 2 to best capture transcriptional adaptations associated with drug-seeking behaviors. All conditions were compared to their respective saline experimental and subregion counterparts to assess the relationship between NAc core and shell subregions across behavioral conditions (Ctx, HC, and Ext). Surprisingly, repetitive context exposure during abstinence (Ctx rats) triggered more robust changes in gene expression (both up- and downregulation) as compared with the other abstinence conditions (HC or Ext; Fig. 2A). Interestingly, this effect was observed predominantly for the NAc core (Fig. 2A). HC and Ext conditions recruited a similar number of DEGs in both NAc subregions, albeit slightly higher in the NAc core in the HC group (or in the shell for Ext) (Fig. 2A).

**Figure 2.**
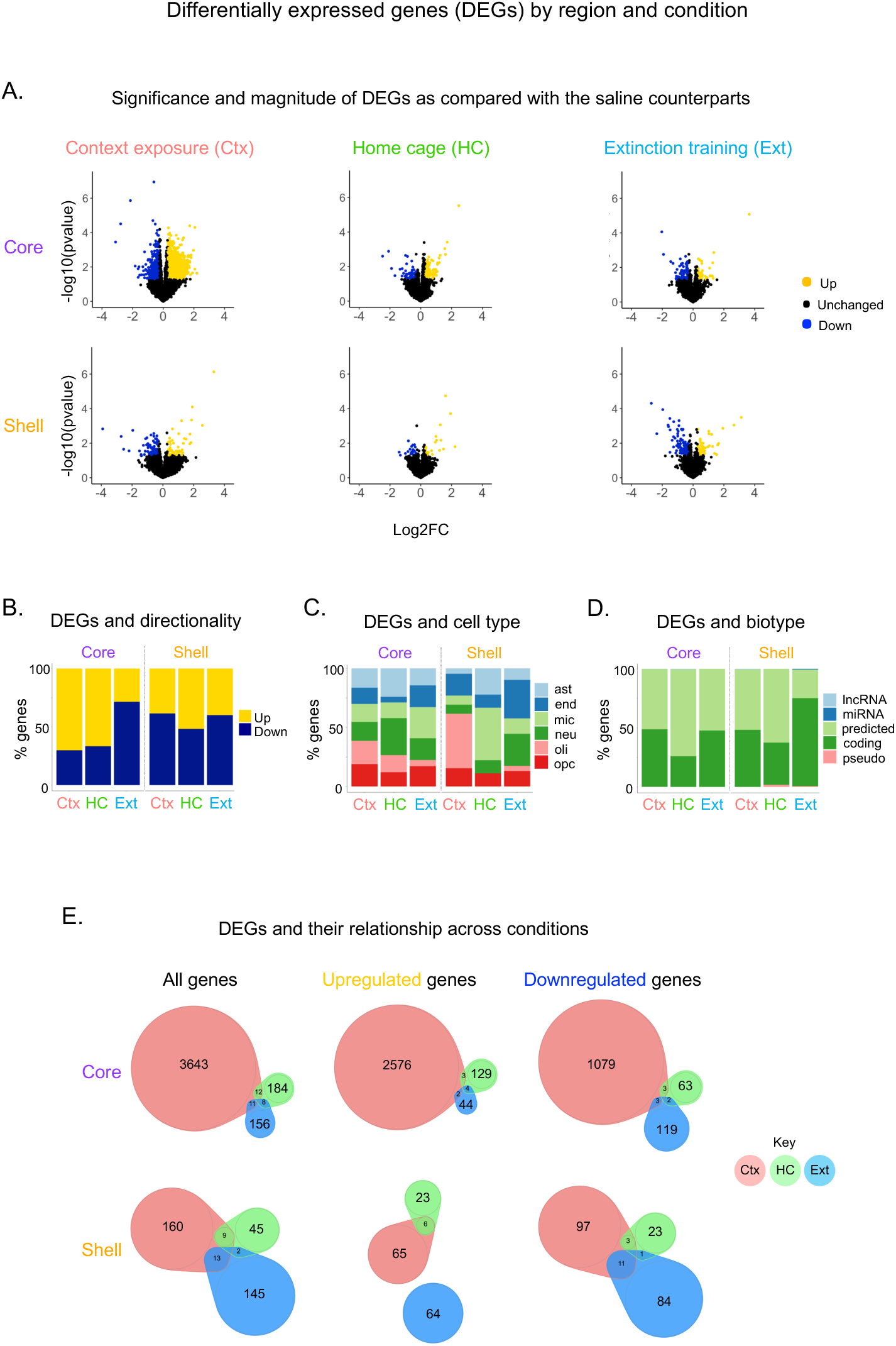
Context-exposed withdrawal disproportionally alters the NAc transcriptome. **A-B.** Volcano plots showing differentially expressed genes (DEGs) from WD/Ext conditions for each NAc subregion. While the Ctx group showed elevated transcriptional changes in the NAc core (particularly), the Ext group displayed more gene expression changes in the NAc shell. Upregulated and downregulated (nominal p<0.05, ± 25%-Fold Change; 1.25) DEGs are represented in yellow or blue, respectively. Distribution (%) and directionality of DEGs. **C-D.** Classification and proportion (enrichment) of DEGs by cell type and biotype presented across experimental groups for each NAc subregion. Cell types: astrocytes (ast), endothelial cells (end), microglia (mic), neurons (neu), oligodendrocytes (oli), or oligodendrocyte precursor cells (OPC). Biotypes: long non-coding RNA (lncRNA), microRNA (miRNA), predicted encoding genes (predicted), protein-coding genes (coding), non-protein-coding genes (pseudogene). **E.** Venn diagrams depicting the proportion of DEGs and their relationship (shared/overlapping) across WD/Ext conditions and NAc subregions.

Further examination revealed more upregulated DEGs in the NAc core for Ctx and HC groups, with an increased number of downregulated DEGs observed in the Ext group (Fig. 2B). A balanced distribution of up- and downregulated DEGs was observed within the NAc shell (Fig. 2B), consistent with the NAc core driving seeking behaviors and the shell more involved in extinction (inhibitory) processes. Cell-type categorization of DEGs across subregions and conditions identified enrichment in five cell types: astrocytes, endothelial cells, microglia, neurons, oligodendrocytes, and oligodendrocyte progenitor cells (OPC). A particular enrichment for astrocytes and neurons was observed in the NAc core HC group and for microglia in the core Ext group, with no enrichment observed for the Ctx group. Within the shell, enrichment for oligodendrocytes, for microglia, and for endothelial cells and neurons was strongly observed in the Ctx, HC, and Ext groups, respectively (Fig. 2C; see also Fig. S2A including non-specific/annotated DEGs). Further biotype analysis showed that the majority of identified DEGs were protein-coding or predicted genes, with few lncRNAs or miRNAs identified (Fig. 2D; see also Fig. S2B for annotated only). This is unsurprising, given that gene annotation focuses largely on protein-coding genes. To determine overlapping and unique DEGs between forced abstinence modalities in the NAc core and shell, intersectional Venn diagrams were generated. Overall, the Venn diagrams illustrate the far greater number of DEGs in the Ctx group, with very little overlap seen across the three abstinence conditions. These findings highlight that distinct clusters of DEGs may drive withdrawal and extinction scenarios in a subregion-dependent manner (Figs. 2E and S2C).

### Genome-wide comparisons of transcriptomic patterns across WD/Ext conditions

Given the lack of converging statistically significant DEGs across abstinence conditions, we next determined if genome-wide patterns of gene expression – analyzed in a threshold-free manner – were similarly divergent by use of stratified rank-rank hypergeometric overlap (two-sided; RRHO2) ^27^. We observed discordant transcriptional patterns between the HC and Ctx conditions in both NAc subregions (Fig. 3A), a surprising finding given the similar levels of drug-seeking behavior exhibited by these two groups. A discordant transcriptional pattern was also observed when comparing Ctx and Ext conditions (Fig. 3A), which suggests distinct molecular adaptations upon context extinction alone vs. full extinction (context/levers/cues). Interestingly, when the HC and Ext groups were compared, concordant patterns within both the NAc core and shell were found, perhaps capturing shared adaptations associated with these two very different abstinence conditions and drug-associated effects. Direct core:shell comparisons within each experimental condition suggest largely concordant patterns within these NAc subregions (Fig. 3B). Together, these transcriptional landscapes unbiasedly support the resulting behavioral profiles after various WD/Ext modalities.

**Figure 3.**
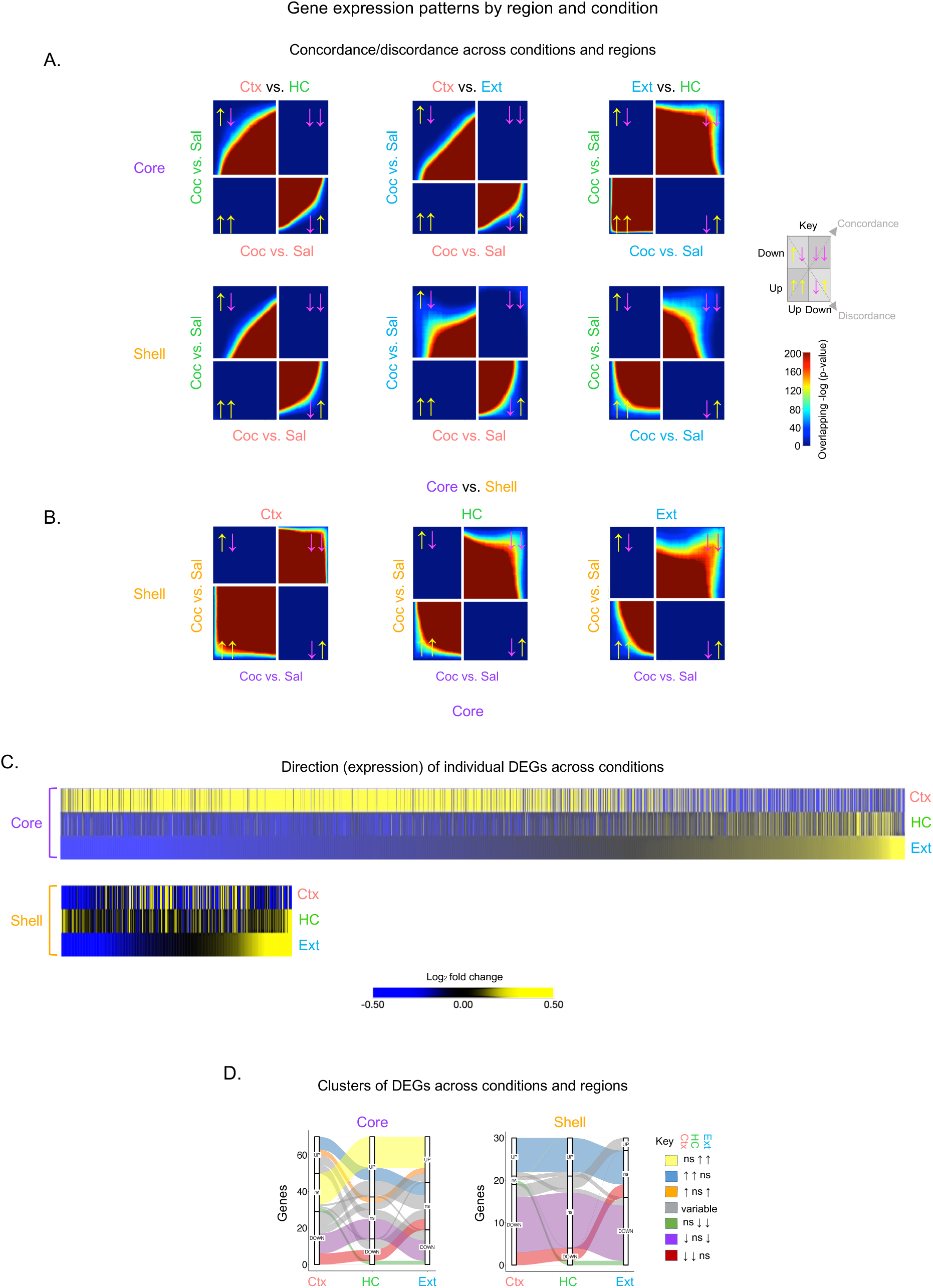
Gene expression patterns across WD/Ext conditions and NAc subregions. **A.** Global threshold-free comparisons of gene expression by Rank-Rank Hypergeometric Overlap (RRHO2) plots showing the degree of transcriptional concordance/discordance across conditions and regions. A strong discordance was observed when the Ctx group was compared with either the HC or Ext group, while HC and Ext displayed transcriptional concordance. Both of these patterns were seen in the NAc core and shell (**B**). Concordance is indicated by the distribution of genes in the bottom left and top right quadrants denoting co-upregulation (yellow arrows in key) and co-downregulation (magenta arrows in key), respectively. Discordance is indicated by the distribution of genes in the top left and bottom right quadrants indicating oppositely regulated genes (up-down; yellow/magenta arrows and down-up; magenta/yellow arrows, respectively). The overlapping intensity is color-coded, where genes are sorted from most to least significant from middle/center to corner in each quadrant. **C.** Union heatmaps generally confirm these RRHO2 patterns by displaying the distribution of all DEGs (upregulated: yellow or downregulated: blue), where the Ctx group displays little overlap with the other two groups. **D.** Alluvial plots highlighting clusters of DEGs with particular expression patterns across conditions and NAc subregions. Plots are stratified by behavioral condition (x-axis) to display the number of DEGs assigned to each pattern/cluster/alluvium (color-coded). Upregulation (↑), downregulation (↓), non-significant (ns), and variable indicate a combination of ↑↓ or ns (no relationship/overlap) for each particular condition.

We next asked whether these threshold-free genome-wide transcriptional patterns are replicated for statistically significant DEGs. Union heatmaps comparing the experimental groups by NAc subregion similarly depicted divergent patterns between the Ctx group and the HC and Ext groups. Also consistent with the RRHO2 analysis, the latter two groups (HC and Ext) shared similar fold change (Log_2_FC) patterns of gene expression. Alluvial plots visualize the flow of DEGs clustered by direction of change that comprise the transcriptional patterns observed across conditions. From this analysis, seven different dynamic clusters were captured that display concordance in at least two conditions (Fig. 3D, all non-gray colors); clusters showing differential, variable patterns across conditions are displayed as gray curves (Fig. 3D). For instance, the yellow cluster (NAc core) demonstrates concordant upregulation in the HC and Ext conditions but not in the Ctx condition, consistent with RRHO2 results. The purple cluster demonstrates concordant downregulation between Ctx and Ext but not HC, also observed in the union heatmaps (Fig. 3C; NAc shell). These data further suggest that distinct transcriptional programs may underlie the neurobehavioral adaptations across the three abstinence conditions.

### Predicted biological functions, upstream regulators, and transcription factors in NAc subregions across WD/Ext conditions

We next bioinformatically assessed key regulatory mechanisms associated with gene regulation by abstinence condition and NAc subregion. We first used Gene Ontology (GO) analysis to link the enrichment of DEGs across WD/Ext conditions and NAc subregions with known biological functions (Fig. 4A). Top GO terms for the NAc core implicated growth factors and extracellular matrix (Ctx) and RNA biosynthesis (HC and Ext). In contrast, the NAc shell was associated with DNA replication and receptor signaling/transactivation (Ctx and HC) and extracellular matrix (Ext). Interestingly, Venn diagrams revealed minimal overlapping of the predicted biological functions across conditions within and between NAc subregions (Figs. 4B and S3A), consistently supporting the idea that distinct molecular mechanisms are at play to maintain distinct behavioral profiles. Examination of the genes within each GO term displayed a balanced up- and downregulation of genes in the NAc core, whereas mostly downregulation was observed for the NAc shell in the Ctx group (Fig. S3B, left panels). The HC condition was strongly associated with gene upregulation for both NAc subregions, whereas the Ext condition was largely associated with gene downregulation (Fig. S3B, middle and right panels).

**Figure 4.**
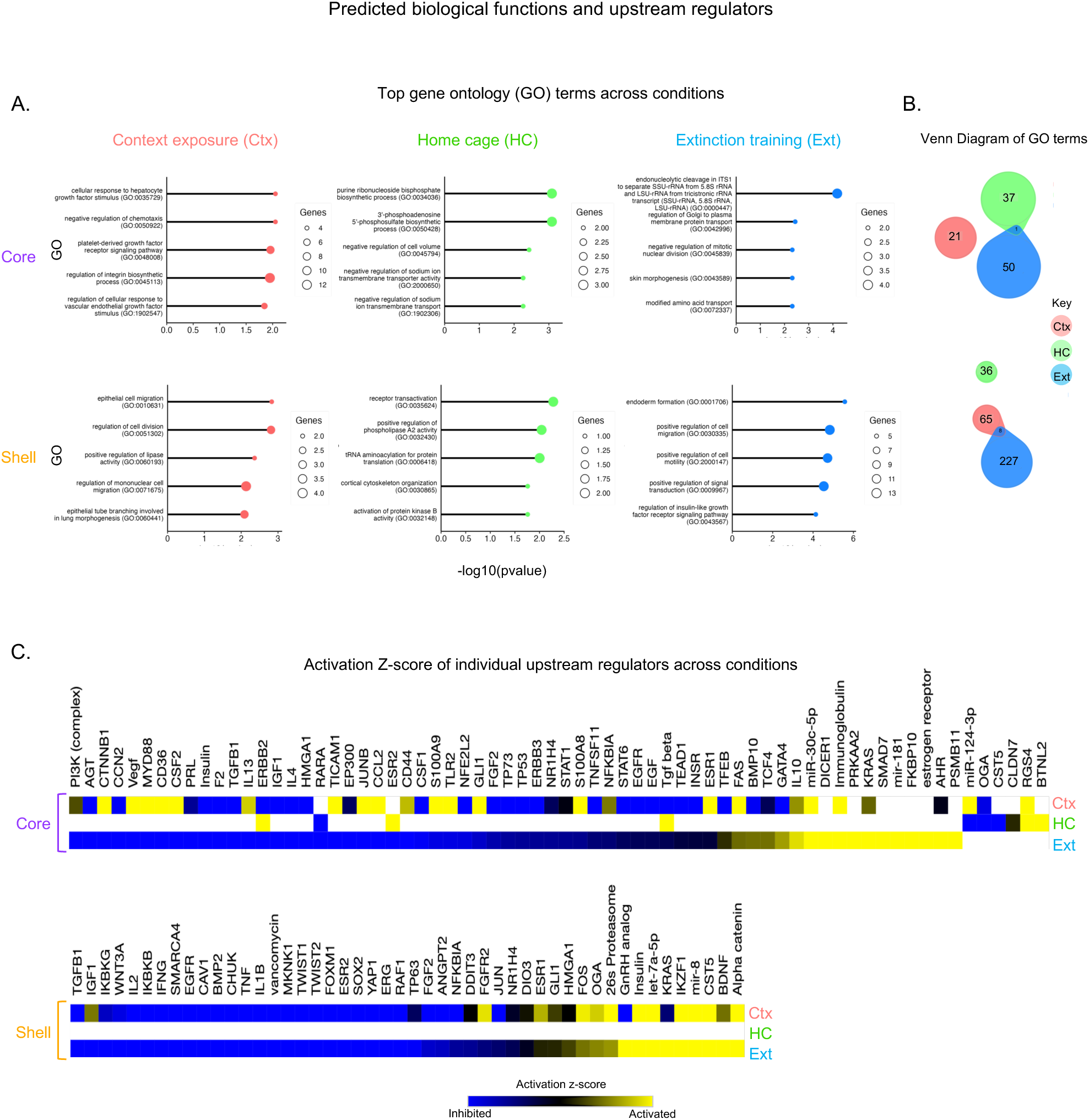
Predicted biological functions and upstream regulators across WD/Ext conditions and NAc subregions. **A.** Top gene ontology (GO) terms in the NAc core featured enrichment of DEGs associated with growth factors, integrins regulation (Ctx), and RNA biosynthesis (HC and Ext), while the NAc shell was implicated in cell division, DNA replication, receptor signaling/transactivation (Ctx and HC) and extracellular matrix dynamics (Ext). **B.** Venn diagrams highlighting different GO terms across conditions with the Ext group involving a higher number of biological functions. **C.** Upstream regulator analysis linked numerous putative regulators to the seeded DEGs with activation or suppressive properties across the WD/Ext conditions in the NAc core or shell. Activation z scores: Activated (yellow) = overrepresentation of targets activated by the regulator; Suppressed (blue) = overrepresentation of targets repressed by the regulator; black/gray = not a predicted upstream regulator.

To further determine molecular mechanisms underlying behavioral adaptations, we performed GO analysis on clustered DEGs from the three behavioral conditions for each NAc subregion (Fig. S4A-B). Six clusters were identified in the NAc core (Fig. S4A), where clusters 1-3 and clusters 5&6 shared similar GO terms, respectively, despite distinct transcriptomic patterns (e.g., extracellular organization and transmembrane transporter activity, respectively). Cluster 4, where DEGs were downregulated in the Ctx group and upregulated in HC and Ext, was associated with dendrite and axon morphogenesis. In the NAc shell (Fig. S4B), 8 clusters were observed, where clusters 1&2, 3-6, and 7&8 each shared similar GO terms (e.g., transmembrane transporter organization, DNA replication, and cytoskeleton assembly, respectively).

To identify regulatory factors that might be responsible for the transcriptional regulation observed, we used Ingenuity Pathway Analysis’s (IPA) Upstream Regulator Analysis. Union heatmaps display regulators found in at least two out of the three abstinence conditions (Fig. 4C, upper: NAc core and lower: NAc shell). Despite similar transcriptional patterns between Ext and HC conditions (Fig. 2), we observed more overlap between Ctx and Ext and few predicted regulators for the HC condition (Fig. 4C). Additionally, within the core, while some regulators were found in common, half (34/65) demonstrated opposing activation status. Given the lack of overlapping statistically significant DEGs, it is perhaps not surprising that different regulators are implicated in the Ctx and Ext groups (Fig. 4C). We identified some previously implicated as well as potential novel regulators. For instance, activation of JUNB and FOS in the Ctx group may support the elevated seeking behavior (Fig. 4C), whereas activation of BDNF in the Ext group could signal extinction learning ^36–39^ (Fig. 4C, lower/NAc shell panel). RARA (retinoic acid receptor α) and RGS4 (regulator of G protein signaling 4) were identified in HC rats and previously associated with cocaine ^40,41^ (Fig. 4C, upper/NAc core panel). Interestingly, this analysis captured changes in estrogen receptors (ESR1/2) including downregulation of these upstream regulators in Ext rats (Fig. 4C, upper/NAc core, and lower/NAc shell panels), where these receptors have been linked to cocaine actions ^42–44^.

We next turned to motif enrichment and discovery analyses (MEME Suite, SEA web-based software) to identify upstream transcription factors predicted to control the regulation of altered genes ^31,45^. The top 30 transcription factor motifs deduced and shared for each behavioral condition demonstrated little overlap between behavioral conditions (Fig. 5A), similar to our Upstream Regulator analysis. Notably, the identified transcription factors were particularly driven by the Ctx condition in the NAc core, suggesting elevated transcriptional regulation. We identified enriched putative motifs for ZNF263 (Ctx group) and RRBE1 (HC group), proteins previously associated with drug exposure ^46,47^ (Fig. 5B, left/middle panels). In the Ext group, a notable enrichment was observed in the NAc shell (Fig. 5A), consistent with the importance of this subregion for extinction learning. Among the top factors, ZNF263 and SP2 were observed, both previously observed in drug conditions ^46,48^ (Fig. 5B, right panels).

**Figure 5.**
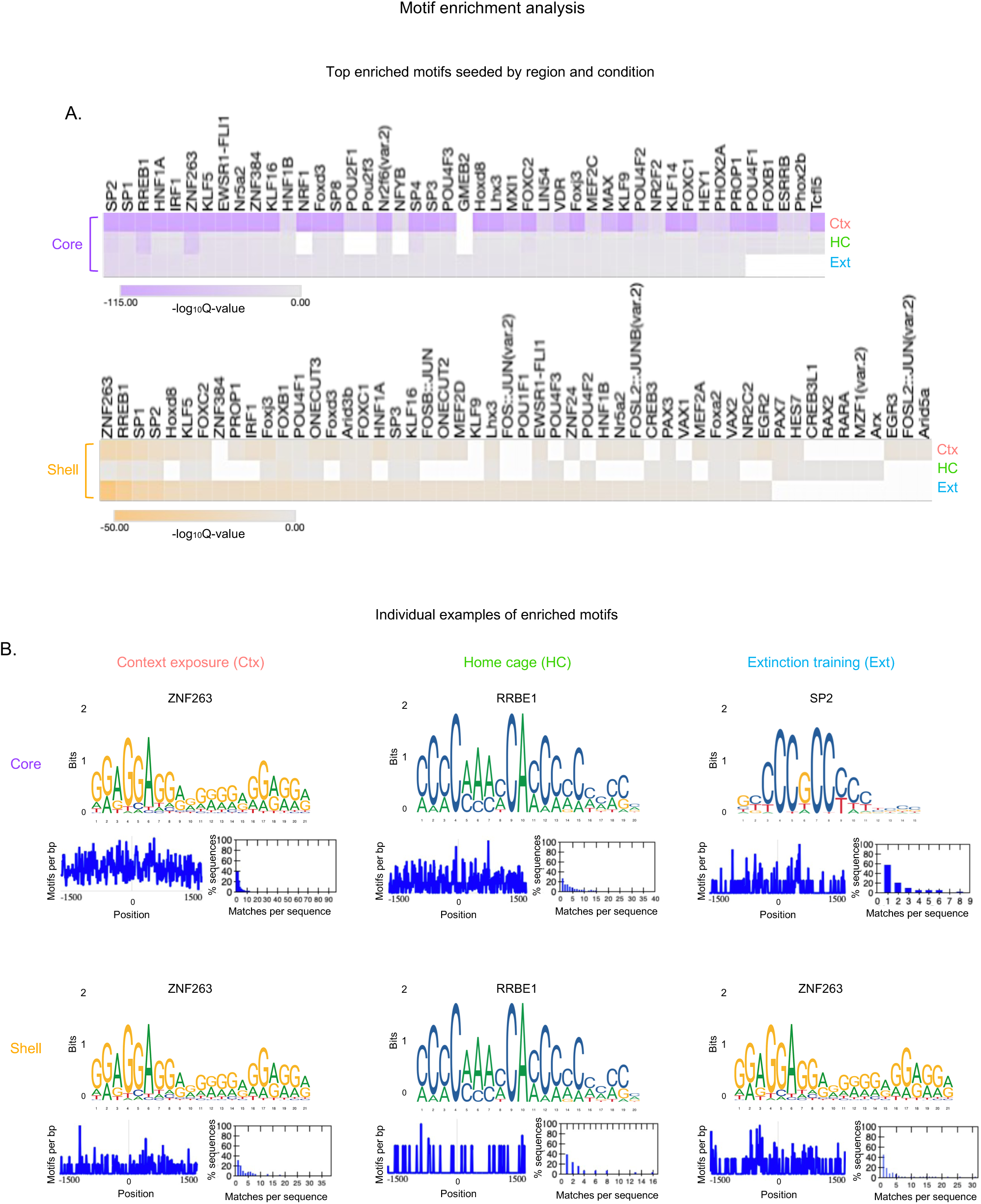
Motif enrichment analysis and identification of DNA binding factors across WD/Ext conditions and NAc subregions. **A.** Top transcription factors deduced across WD/Ext conditions and NAc subregions obtained by Simple Enrichment Analysis (SEA) of DEGs. The identified motifs were overwhelmingly linked to putative factors in the Ctx group within the NAc core, while elevated incidences in the NAc shell were observed for the Ext group. The Q-value represents the minimum false discovery rate (FDR) required to consider a motif significant. **B.** Representation of selected (top) individual motifs (logos, positions, and incidence/matches) and their binding factors. Positional distribution of the best match to the motif in the primary sequences. The dotted line indicated the position of the sequences aligned to the left/right ends or centers. The histograms show the number of motif matches in the primary sequence with at least one match.

### Divergent gene co-expression networks across NAc subregions and WD/Ext conditions

We used MEGENA to determine groups of co-regulated genes across conditions and identified organized hierarchical gene modules (sunburst plots; Fig. 6A). This analysis revealed minimal overlap of gene modules when the Ctx group is compared to either the HC or Ext group (Fig. 6A, NAc core, left/middle panels). By contrast, several gene modules displayed overlap between the HC and Ext groups (Fig. 6A, NAc core, right panel), supporting the uniqueness of the Ctx group with other analyses. Similar, although weaker in magnitude, results were observed under saline conditions (Fig. S5A). In contrast, the NAc shell displayed an overlap of modules between the Ctx group and the HC and Ext groups (Fig. 6A, NAc shell, lower panels), further suggesting subregion-dependent patterns of regulation. The NAc shell showed little overlap under saline groups, suggesting cocaine-specific effects regardless of the abstinence behavioral paradigm. A small number of modules exhibited overlap for each WD/Ext condition for the NAc core and shell (Fig. S6A-B).

**Figure 6.**
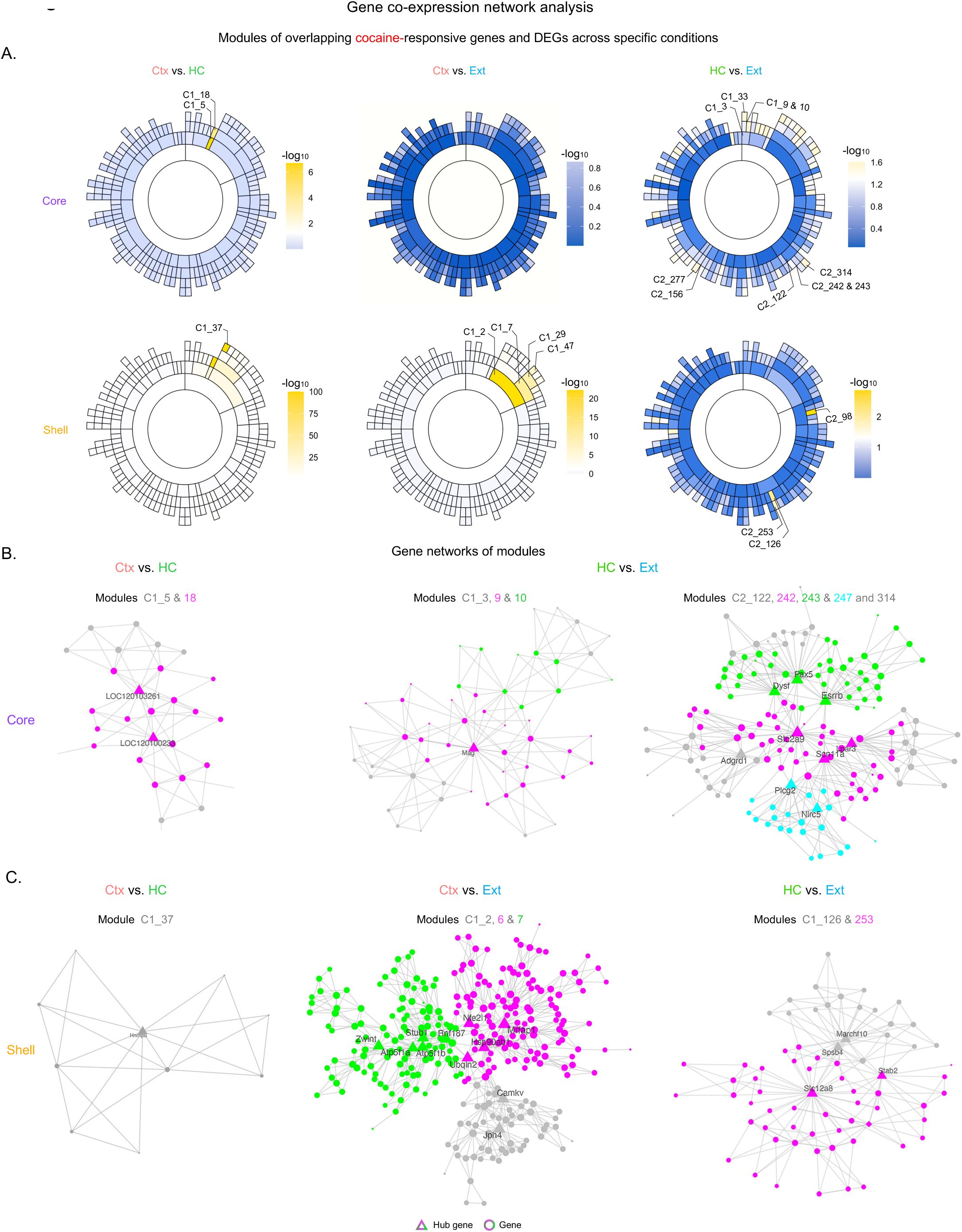
Co-expression network analysis across WD/Ext conditions and NAc subregions. **A.** Sunburst plots, depicting gene co-expression networks by MEGENA, show little overlap (white/blue modules) when the Ctx group is compared with the HC or Ext groups (core/shell; left and middle panels). Still, the presence of overlapping gene modules was observed (yellow modules). Comparison between HC and Ext conditions revealed several shared gene modules (yellow), particularly in the NAc core (top-right panel). Inner and outer rings of the sunburst plots are referred to as parent and child modules, respectively. **B.** ARACNE plots showing gene network and connectivity within some of the overlapping modules across WD/Ext conditions and NAc subregions. These networks also highlight hub/driver and other highly connected genes (π= hub gene; λ= gene).

We used ARACNE to identify and infer gene regulatory interactions and bonafide targets mediated by transcription factor binding and reconstruct the associated networks from our dataset (Fig. 6B-C). Several genes were identified as overlapping hub genes across abstinence conditions. For instance, in the NAc core (Fig. 6B and S7A), small nucleolar RNAs (e.g., “LOC genes”; parent module C1_5) were central to Ctx vs. HC groups (Fig. 6B, left panel), whereas myelin-associated glycoprotein (*Mag*; parent module C1_3) and several others hub genes including *Scn11A* (encoding a sodium channel subunit), *Esrrb* (encoding an estrogen receptor), *Plcg2* (encoding a phospholipase Cγ) (parent module C2_122), and *St8sia6* (encoding a glycosyltransferase) were central to HC vs. Ext groups (Fig. 6B, middle/left panels and Fig. S7A). As for the NAc shell (Fig. 6C and S7B), nuclear ribonucleoprotein (*Hnrnpu*; parent module C1_37) was central to Ctx and HC groups (Fig. 6C, left panel), whereas calcium/calmodulin-dependent protein kinases (*Camkv*, parent module C1_2), heat-shock proteins (*Hsp90ab1*; child module C1_6) and ATP synthesis (*Apt5f1a/b*; child module C1_7) were among the hub genes relevant for the Ctx and Ext groups (Fig. 6C, middle panel and Fig. S7B, left panel). In the HC and Ext groups, genes involved in cell membrane dynamics (membrane-associated ring finger; *March10*; module C1_126), cation-chloride transport (*Slc12a8*; module C1_253), and other unspecified genes (e.g., LOC genes; module C1_98) (Fig. 6C, right panel and Fig. S7B, right panel). Several of these hub genes (e.g., *Camkv, Hnrnpu,* and *Mag*) have been also associated with cocaine exposure ^49–55^.

### Comparison of transcriptional regulation in rat NAc across WD/Ext conditions to transcriptional abnormalities in the NAc of human CUD

We recently demonstrated convergence between gene expression changes seen in NAc after cocaine self-administration in mouse and CUD in humans (Mews et al., 2023), but such convergence has never been examined across WD/Ext conditions. We, therefore, proceeded to compare global transcriptomic expression patterns, DEG distribution, and GO pathway analysis of Ctx, HC, or Ext groups in rats to CUD subjects who died from cocaine overdose (Fig. 7A). RRHO2s revealed a global concordance between Ctx and CUD in both NAc core and shell, whereas HC and Ext showed general discordance compared with CUD (Fig. 7B). These patterns were also observed when comparing statistically significant DEGs by NAc subregion (Fig. 7C, core and shell). However, a notable overlap in downregulated genes was observed between HC, Ext, and CUD conditions that were not observed within the appropriate RRHO2 quadrant (Fig. 7C, core and shell). Highlighting convergence between Ctx and CUD, particularly within the NAc core, Venn diagrams revealed a subset of DEGs shared between Ctx and CUD variables (Fig. 7D), with several GO terms represented (e.g, phospholipid processes, histone acetylation, and calcium signaling) (Table 1). Indeed, several of these genes (e.g, *SMAD3*, *HTR7*, *PPP1R17*) ^56–60^ and their upstream regulators (e.g., STAT3, AR, mir-124) have previously been associated with cocaine exposure ^61–64^ (Fig. 7E). Together, these results suggest that the Ctx condition especially mirrors transcriptional patterns observed in human CUD.

**Figure 7.**
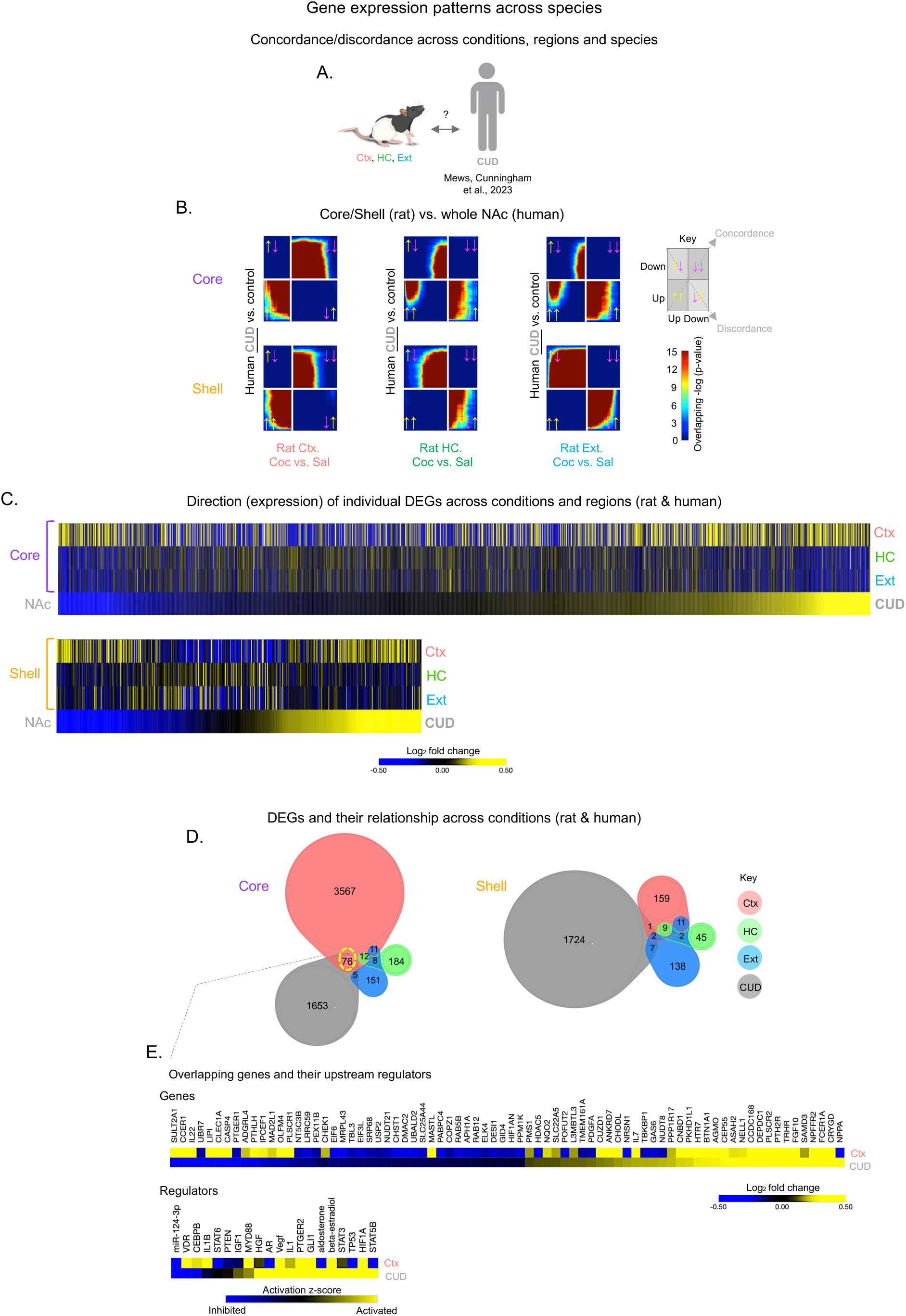
Convergent transcriptional patterns in NAc between Ctx withdrawal in rats and human CUD. **A.** Schematic posing the comparison between WD/Ext modalities in rats and human CUD. **B.** RRHO2 plots reveal a robust transcriptional concordance between Ctx rats and CUD (left panels core/shell) and predominant discordant patterns when CUD is compared against HC or Ext groups (middle and right panels core/shell). Concordance/ Discordance; same or opposite direction of yellow /magenta arrows. **C.** Union heatmaps showing a similar distribution of DEGs between CUD and Ctx groups, which is consistent with the RRHO2 results. Upregulation: yellow or downregulation: blue. **D.** Venn diagrams showing the proportion of DEGs and their relationship (overlap) between CUD and WD/Ext conditions in the NAc subregions. Note that the CUD data were obtained from the whole NAc encompassing both subregions (Mews et al., 2023). **E.** Union heatmaps (upper panel) highlighting the shared genes and their expression direction (up = yellow, blue = down) between CUD and Ctx datasets as well as the consequent upstream regulators for the NAc core. Activation z scores: Activated (yellow) = overrepresentation of targets activated by the regulator; Suppressed (blue) = overrepresentation of targets repressed by the regulator; black/gray = not a predicted upstream regulator.

## Discussion

In this study, we used cocaine SA in rats to unveil behavioral and transcriptional patterns of forced abstinence scenarios involving distinct withdrawal or extinction modalities. We identified numerous transcriptome-wide changes in the core and shell subregions of the NAc that associate with the addictive continuum in rats that experienced cocaine withdrawal in their home-cage or in the previous drug context, or experienced full extinction conditions. Characteristic behavioral outcomes were revealed and linked to distinct gene expression patterns. Of particular interest was the observation of enhanced drug seeking in rats undergoing context withdrawal, whose transcriptional profiles converged with those of humans with CUD.

### Forced abstinence modalities

Withdrawal and extinction phases have been classically modeled and studied in animals for decades ^18,65–68^. These strategies have been largely centered on applying forced abstinence either in home-cages or under full extinction conditions. This is commonly followed by re-exposure to drug contexts and cues to measure drug-seeking behavior ^65,69^. Full extinction training involves dissociating the prior drug context, levers, and cue lights from drug access. However, a partial form of extinction is also possible, wherein animals are exposed repeatedly to the previous drug context, but with the omission of other drug-associated triggers (e.g., levers and cue lights). Here, we directly compared all three of these preclinical models.

While contextual extinction during fear conditioning has been found to reduce fear responses ^70–72^, we observed the opposite with the Ctx group. This procedure led to an increase in drug-seeking behavior to the level or greater than that of animals experiencing home-cage abstinence. It is possible that this repeated contextual exposure increased drug-seeking behavior by potentiating the incubation of craving as has been observed during cue/context-induced relapse ^73,74^. Another possibility is that drug contexts may act as “occasion setters” ^75,76^ and therefore not be equally valued compared to the drug-associated instruments (levers) and cues that are directly linked to the drug itself ^77,78^. This is very different from conditioned place preference (CPP) scenarios, where the context represents the only reference associated with the drug ^79,80^. Thus, repeated omission of instruments and cues in the original context could lead to increasing the expectation of obtaining the drug.

Full extinction training, on the other hand, impacts these variants by simultaneously extinguishing drug-paired cues and instruments in the same context, thereby promoting learning processes against all potential drug stimuli ^9,81^. Still, a key challenge is knowing whether the animals refrain from drug seeking because they learned the lack of drug availability or because they developed a reduced drug craving and motivation per se. Consistent with our observed increase in drug seeking after context exposure in rats, this intervention has shown limited clinical success ^82^. Subjects commonly develop compulsive drug taking in a given context with cues, while learn extinction in a different setting, and they often relapse upon returning to the original context and cues ^83^. To our knowledge, this is the first study to directly compare these three WD/Ext modalities in male rats. Future research is needed to determine whether similar patterns hold for females and under other context-associated paradigms (e.g., renewal; ABA).

### Transcriptional programs in NAc under forced abstinence modalities

We show that the three WD/Ext procedures associate with highly distinct transcriptomic patterns in the NAc core and shell, which shed new light on the neurobiology of CUD. Notably, repeated context exposure during withdrawal induced the strongest transcriptional response in this brain region. This transcriptional burden was supported by the upregulation of genes in the NAc core particularly, which were largely protein-coding genes from multiple cell types. This was accompanied by biological functions involved in sterol (lipid) metabolism and membrane (integrin, receptors, lipids) regulation, processes that are highly impacted in drug addiction ^84–88^, as well as upregulation of key upstream regulators, such as JUNB, estrogen receptors (core) and FOS (shell), that are also implicated in addiction ^89,90^. Motif analysis revealed singular enrichment for these transcription factors in core and shell subregions. Interestingly, while these results largely differed from transcriptional landscapes of animals receiving full extinction training, these results also diverged from those animals experiencing home-cage withdrawal. This discordance vs. full extinction suggests that extinction training may counteract some of the key features of withdrawal and relapse at the behavioral and transcriptional levels ^37^. However, the unexpected transcriptional discordance between context and home-cage withdrawal, even when both conditions associate with robust drug-seeking behavior, suggests that different biological pathways lead to relapse and that the withdrawal context matters ^14,78,81,83^. Stated another way, our results indicate that different withdrawal paradigms may trigger similar relapse responses by recruiting different molecular and cellular mechanisms. These findings inform future individualized medicine approaches for patients with CUD for whom relapse may be triggered by different paraphernalia and settings ^7,91,92^.

A surprising finding was the observation of similar gene expression patterns between animals receiving home-cage withdrawal and full extinction training. Such transcriptional convergence between these two conditions could represent shared withdrawal-associated changes in the NAc despite the extinction training. Nonetheless, a close look at the distribution of statistically significant DEGs, upstream regulators, and biological functions highlights clear transcriptomic differences for the NAc core and shell between these two conditions, as previously suggested ^15,17,59,93^. Some of these differences could reflect contextual withdrawal differences, but importantly we incorporated corresponding saline controls that account for the context, cues, and lever-associated stimuli independent of the drug in this study.

### Gene co-expression networks in NAc subregions across WD/Ext conditions

Transcriptomic differences between NAc core and shell during abstinence have been demonstrated previously but with limited characterization ^17,93,94^. In this study, RRHO2 analyses – which are threshold-free – revealed concordant gene expression patterns between NAc core and shell, whereas differential gene expression analyses revealed pointed differences among our behavioral scenarios throughout the study. For instance, Venn diagrams displayed very little overlap between core and shell in the distribution of DEGs (Fig. S2C) for each behavioral condition, an effect that was also observed in gene ontology analyses (Fig. S3A). Our search for hub genes or master transcriptional regulators also showed differences between these subregions. For example, the upstream regulator BDNF was upregulated following extinction in the NAc shell, but downregulated in animals receiving contextual withdrawal, in agreement with other studies suggesting that BDNF in the shell is a key molecule driving extinction learning ^37–39,95^. Motif analyses also display enrichment of zinc finger proteins, BDNF signaling (e.g., with CREB being downstream of BDNF), MAP kinase (MAPK) signaling (e.g., RRBE1), and specific transcription factors like FOX and FOS/JUN proteins, all relevant for drug-related behaviors and plasticity ^96–98^. Lastly, network analysis revealed minimum overlap between gene co-expression modules for NAc core and shell, opening a new way to search for unique hub genes involved in each subregion across WD/Ext conditions.

Only limited reports have examined the transcriptional signatures and interplay across withdrawal and extinction modalities within the NAc core and shell. Thus, new rodent and human studies will be needed to understand the molecular features in the brain’s reward circuitry under these various conditions. However, several hub genes associated with small nucleolar RNA (including several LOC genes) and ribonucleoproteins (*Hnrnpu*) that are key in RNA processing (e.g., methylation, uridylation, and stabilization) stand out when Ctx and HC groups are compared, highlighting withdrawal features as opposed to extinction ^49,54^. Gene co-expression between Ctx and Ext reveals cell signaling (*Camkv*) and metabolism processes (*Atp5f*), perhaps critical to instrumental or contextual WD/Ext learning ^51,52,55^. Notable overlapping results between HC and Ext groups identified several likely regulatory (hub) genes involved in ion transporters and channels (*Slc12a8/9, Scn11a*), estrogen actions (*Esrrb*), and myelin maintenance (*Mag*), potentially signaling spontaneous extinction- or withdrawal-associated processes of cocaine ^42–44,50,53,99–102^.

### Transcriptional convergence in the NAc between rat WD/Ext and human CUD

Capitalizing on recent human studies, we integrated the present rat transcriptional dataset with the NAc transcriptomes from individuals with CUD who died from cocaine overdose ^19^. This integration enabled us to identify gene signatures that are translationally significant for understanding CUD. Notably, animals receiving Ctx withdrawal showed a strong convergence in their global transcriptome with that of CUD, characterized by shared DEGs and common upstream regulators. Examples of these shared DEGs include *HTR7, FGF10, SAMD3, NPFFR2, and PABPC4*, whereas GLI1, STAT3, estradiol, and mir-124-3p were among the overlapping upstream regulators. Conversely, HC withdrawal and full extinction conditions largely showed significant divergence from the CUD transcriptome, a discrepancy that may mirror the specific addiction stage when the CUD subjects died. Although home-cage abstinence is the gold standard approach to studying the neurobiology of drug withdrawal in rodents, the fact that the Ctx group converges the most with the human CUD data suggests especially strong validity of this particular form of abstinence in modeling key aspects of cocaine withdrawal in humans. Moreover, the fact that full extinction associates with unique gene expression patterns in the NAc supports the strategy of using extinction-based approaches as a preferred non-invasive treatment for CUD ^103^. Future research should aim to dissect the transcriptional mechanisms (e.g., cell specificity and epigenetic regulation) underlying HC withdrawal, Ctx withdrawal, and full extinction to pinpoint gene expression changes linked to specific behavioral outcomes of these states. Examining different time points and gene expression after relapse or context renewal, as well as extending this study to additional brain reward regions, will be crucial next steps.

## Conclusions

As molecular biological and bioinformatic analyses progress, the use of these preclinical approaches is fundamental to modeling some of the most devastating features of SUDs. The transcriptome-wide mapping of gene expression changes in the rat NAc core and shell induced by three very different abstinence conditions provides novel biological insight into CUD. Our data provide a comprehensive assessment of individual genes, gene networks, biological functions, upstream regulators, and other molecular properties observed in the NAc during withdrawal vs. extinction. Our findings also highlight how transcriptomic profiling might be used to formulate and perfect new behavioral strategies (e.g., extinction-based training) to counteract specific aspects of withdrawal and relapse for CUD.

## Acknowledgments

This work was supported by NIH Grants: 5T32DA007135-33 & K01DA054306 to FJMR. R01DA007359 & P01DA047233 to EJN. R00AA027839 to PM and F99NS129172 to AMT. The authors want to thank Joseph Landry and James Callens (Yasmin Hurd Lab); and Giselle Rojas, Kyra Schmidt, Nathalia Pulido, Katherine Beach, Stephen Pirpinias, Angélica Torres-Berrío, Yun Young Yim, Clementine Blaschke, Kinneret Rosen, and Ezekiell Mouzon (Nestler Lab), for technical support and scientific feedback. We also want to thank Drs. Paul Kenny and Yavin Shaham for their advice and scientific discussions.

## Supplementary Figure Legends

**Figure S1 (for Fig. 1).**
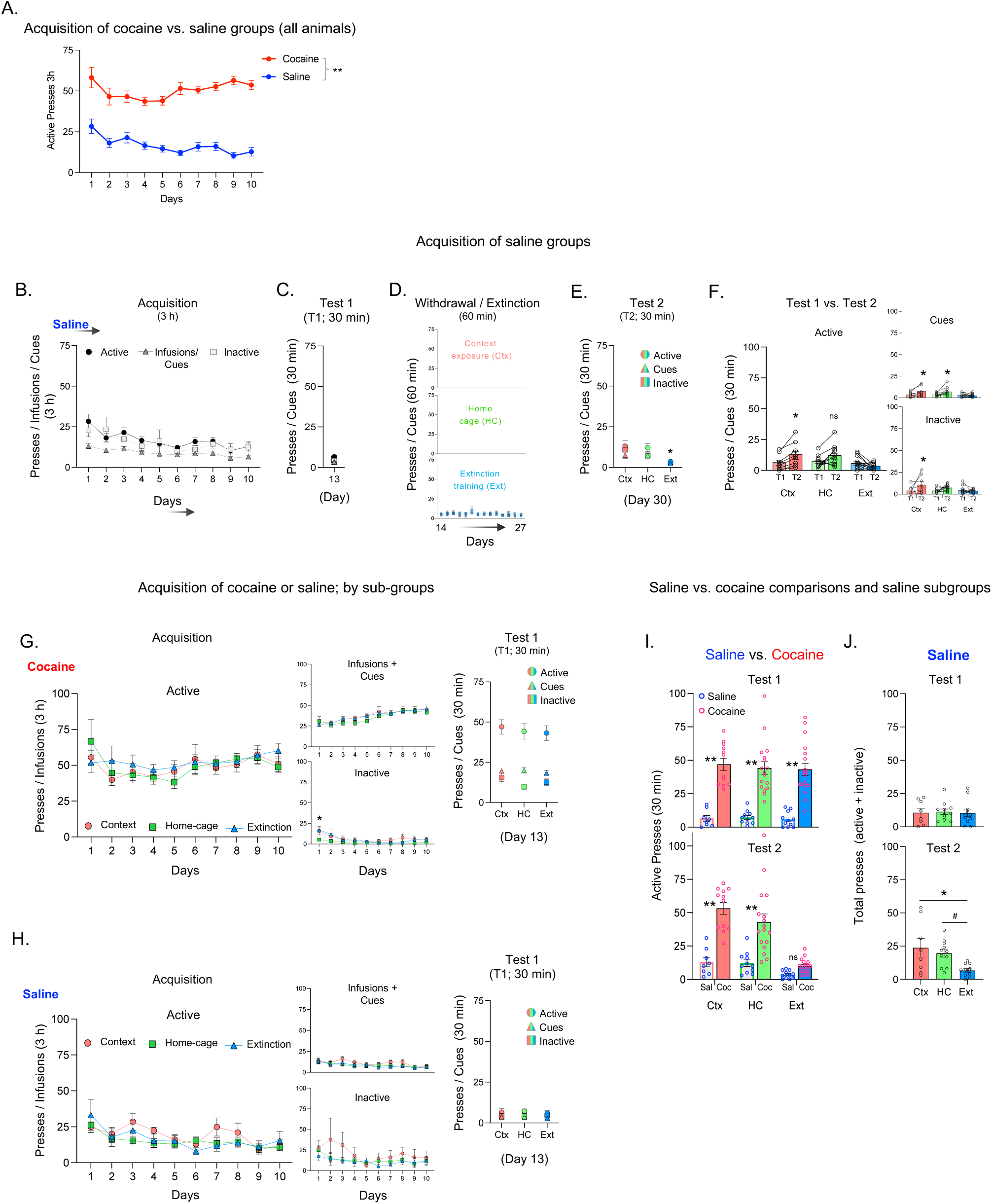
Comparisons between saline- and cocaine-treated WD/Ext subgroups. **A.** Acquisition of cocaine vs. saline SA. **B-F.** WD/Ext protocol/design in saline-treated animals. **G-H.** Acquisition and Test 1 of WD/Ext groups in saline- or cocaine-SA animals. **I.** Comparing the level of seeking behavior (active presses) between cocaine and their corresponding saline subgroups in Tests 1 and 2 (T1 and T2). **J.** Total lever presses (active and inactive) in saline-treated groups during Tests 1 and 2. Saline subgroups: Saline, Ctx: n=8; HC: n=11; Ext: n=11. All data are shown as mean ± SEM. **p≤0.01, *p≤0.05, ^#^p≤0.10.

**Figure S2 (for Fig. 2).**
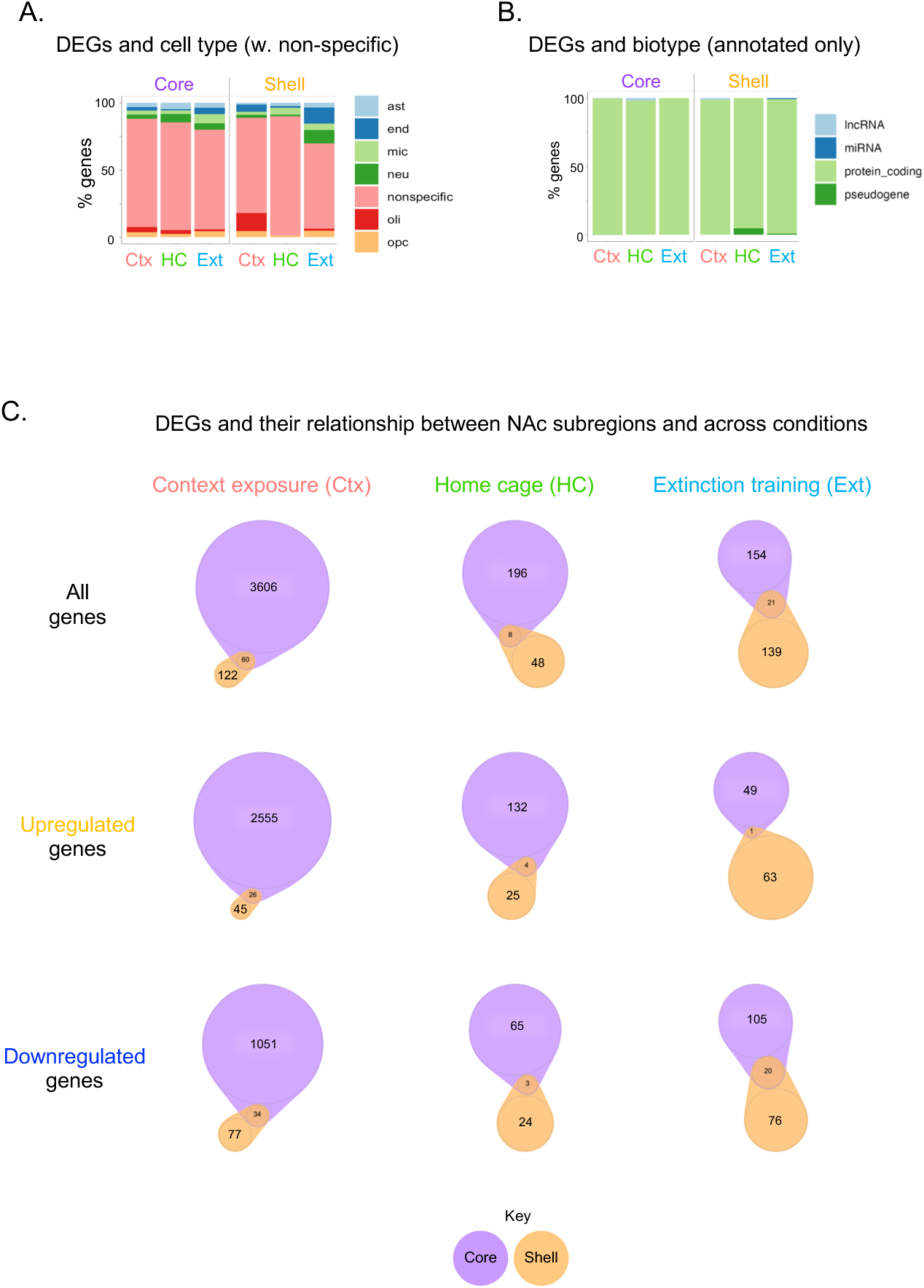
Classification of DEGs and NAc core vs. shell comparisons. **A-B.** Classification and proportion (enrichment) of DEGs by cell type (including non-specific cell classification) and biotype (annotated only) were presented across experimental groups for each NAc subregion. Cell types: astrocytes (ast), endothelial cells (end), microglia (mic), neurons (neu), oligodendrocytes (oli), or oligodendrocyte precursor cells (OPC). Biotypes: long non-coding RNA (lncRNA), microRNA (miRNA), predicted encoding genes (predicted), protein-coding genes (coding), non-protein-coding genes (pseudogene). **C.** Venn diagrams depicting the proportion of DEGs and their very limited overlap when NAc core and shell are directly compared across WD/Ext conditions.

**Figure S3 (for Fig. 4).**
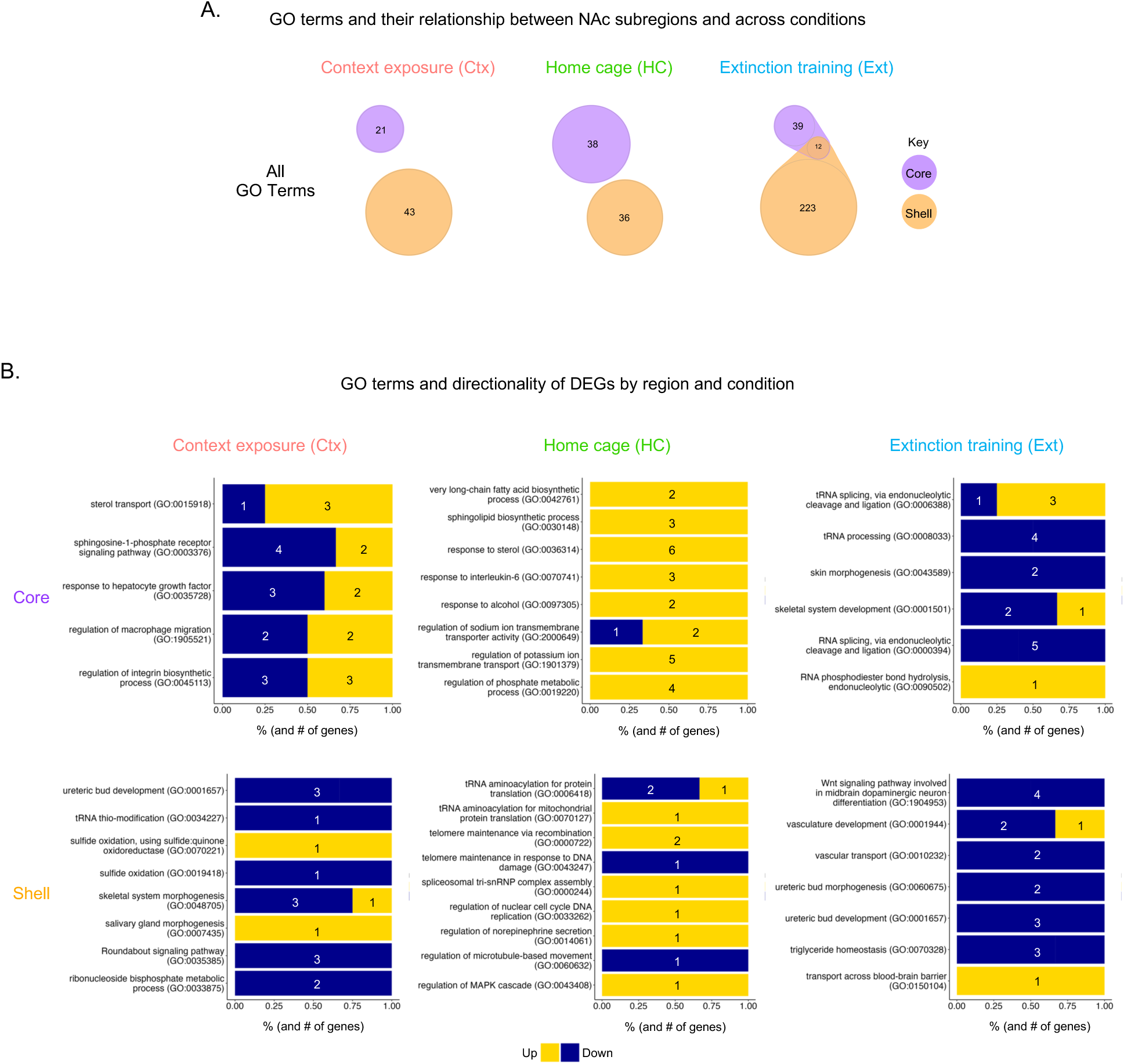
Predicted GO term analyses and NAc core vs. shell comparisons. **A.** Venn diagrams highlighting different GO terms across conditions with little overlap between NAc core and shell. **B.** GO term enrichment and directionality of DEGs across WD/Ext conditions and NAc subregions. Numbers inside the bars represent the number of genes. Yellow = upregulation; Blue = downregulation.

**Figure S4 (for Fig. 3 & 4).**
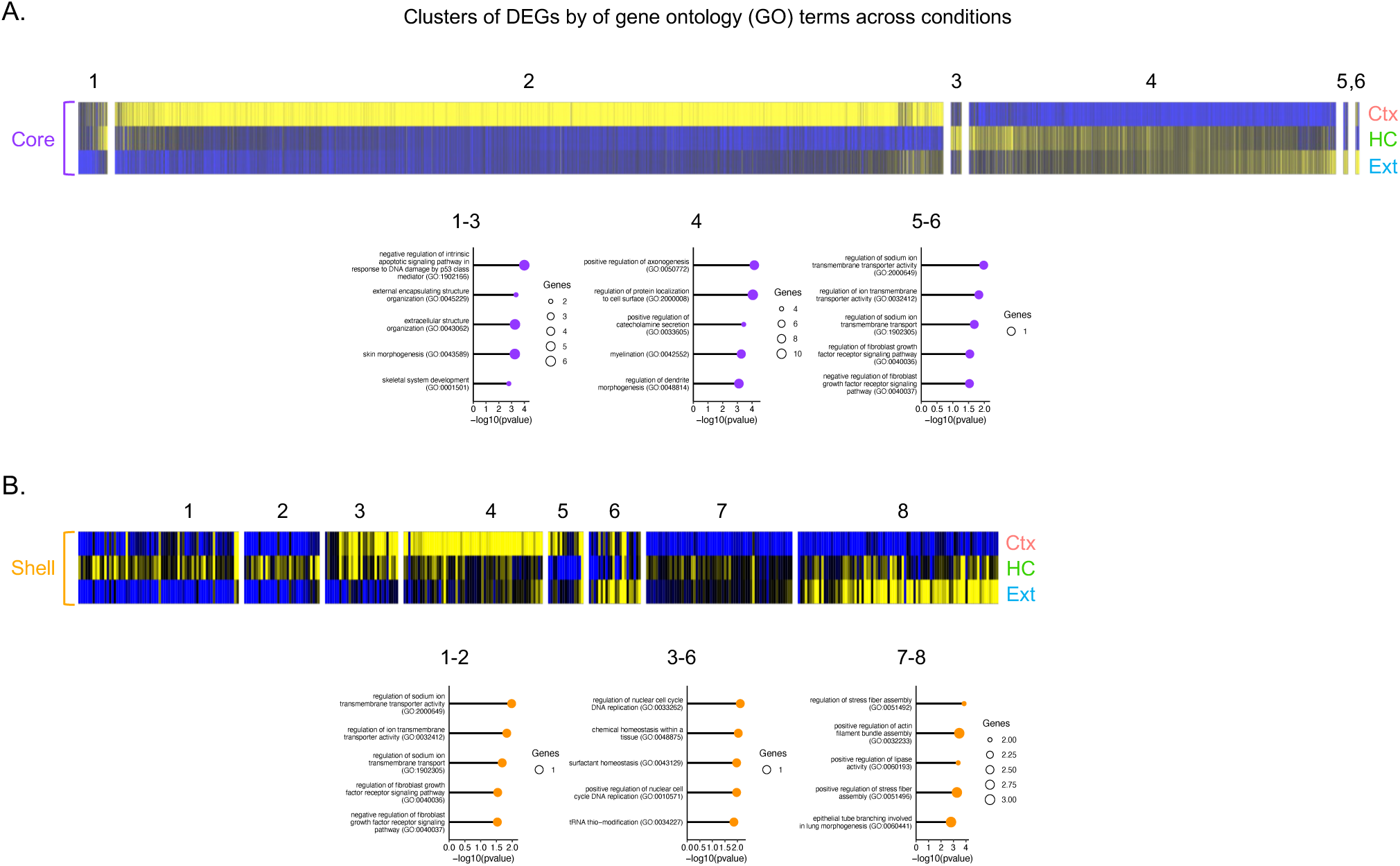
Cluster analyses and predicted GO terms. **A-B.** Union heatmaps from NAc core and shell grouped by shared GO terms (bubble plots). Yellow = upregulation; Blue = downregulation.

**Figure S5 (for Figs. 6).**
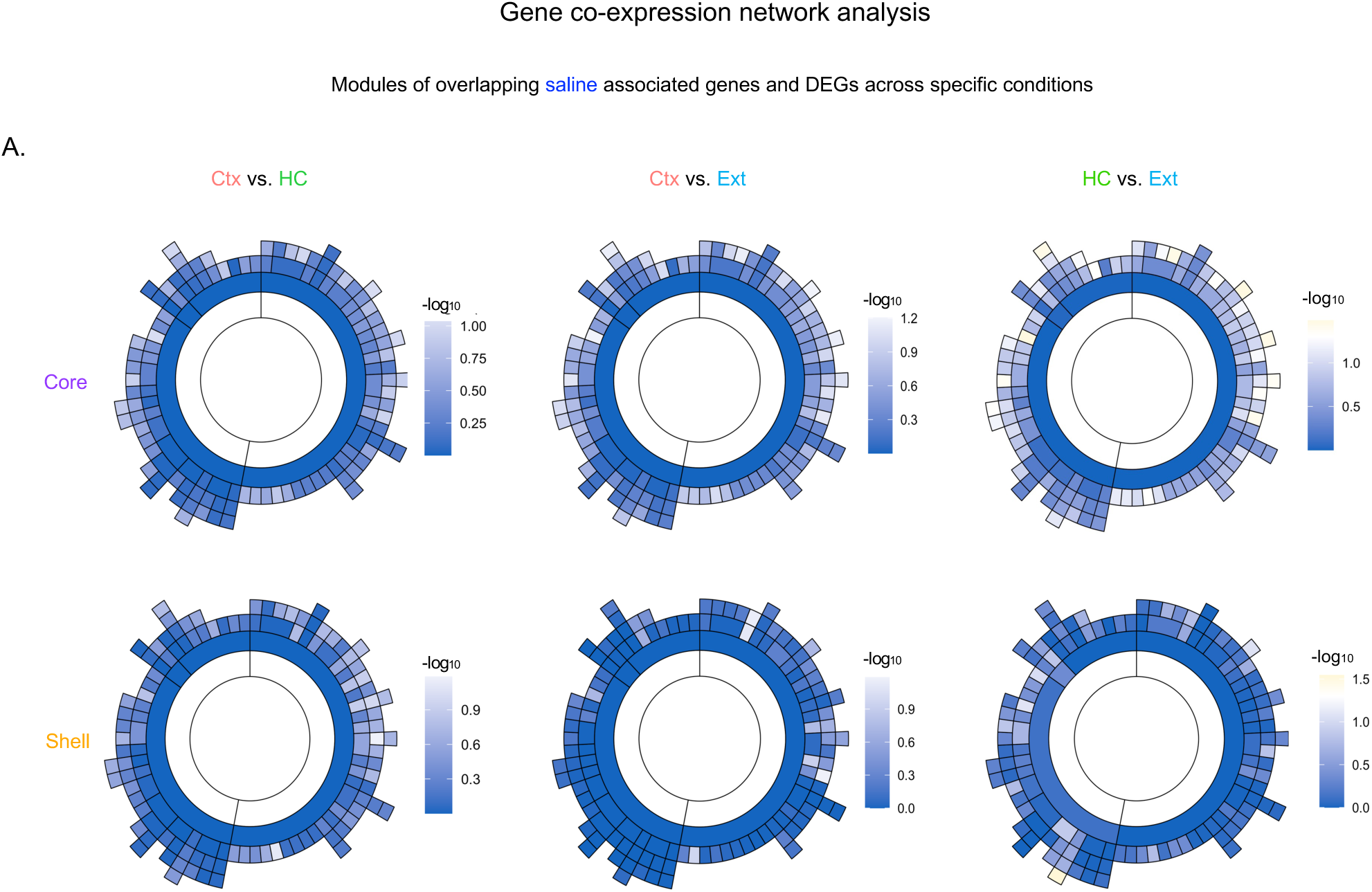
Co-expression network analysis of saline-treated groups for each NAc subregion. **A.** Sunburst plots showing little overlap (white/blue modules) when the Ctx group is compared with the HC or Ext groups (core/shell; left and middle panels). Comparison between HC and Ext conditions revealed a few overlapping modules (yellow; right panels). Inner and outer rings of the sunburst plots are referred to as parent and child modules, respectively.

**Figure S6 (for Fig. 6).**
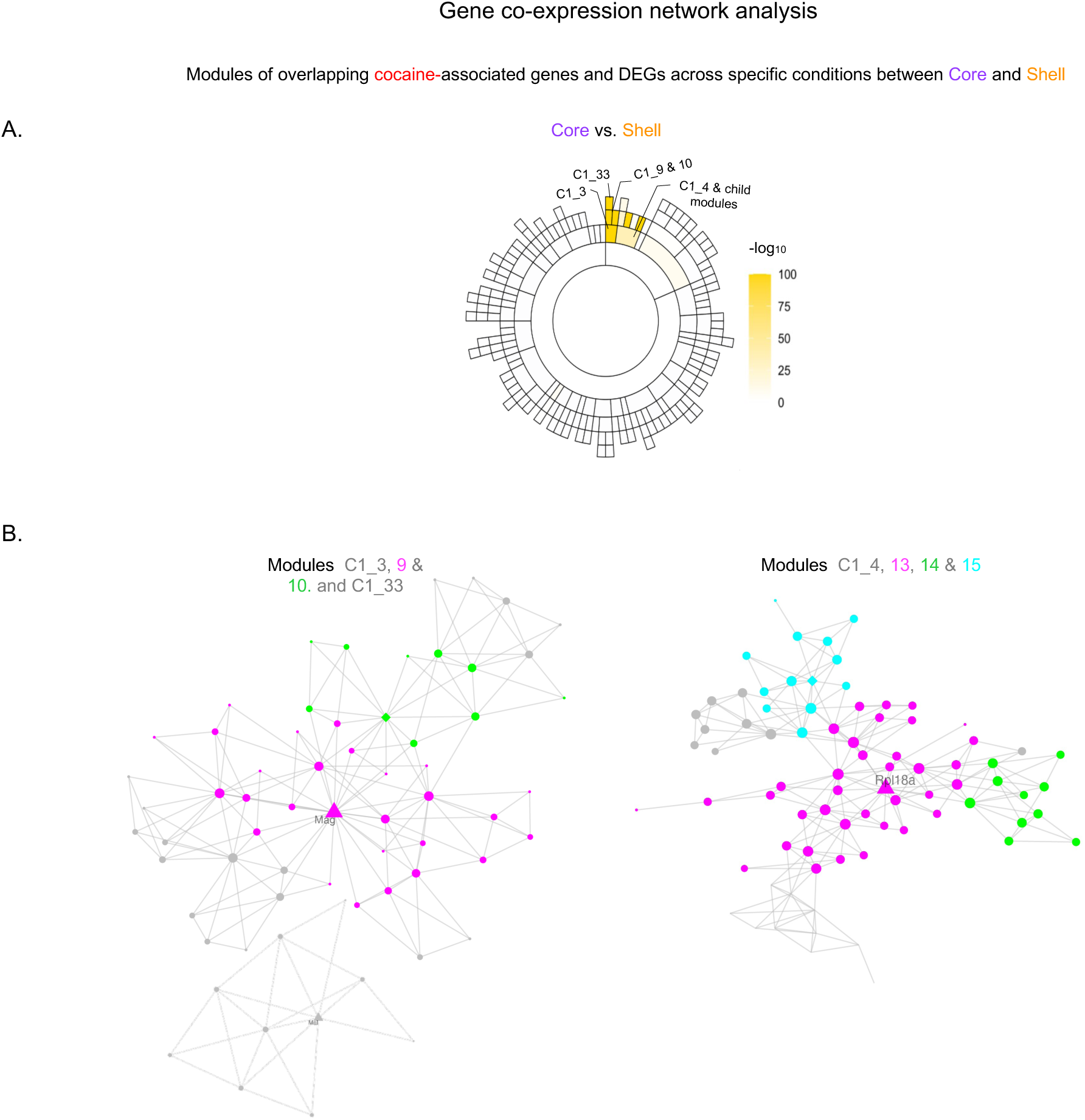
Co-expression network analysis of NAc core vs shell. **A.** Sunburst plots showing little overlap (white/blue modules) between NAc core and shell. Overlapping modules are highlighted in yellow. Inner and outer rings of the sunburst plots are referred to as parent and child modules, respectively. **B.** ARACNE plots showing gene network and connectivity within some of the overlapping modules between NAc core and shell. These networks also highlight hub/driver and other highly connected genes (p = hub gene; l = gene). *Rpl18a* = ribosomal protein L18A; *Mag* = myelin-associated glycoprotein; Mal = Myelin and lymphocyte protein (also known as T cell differentiation protein).

**Figure S7 (for Fig. 6).**
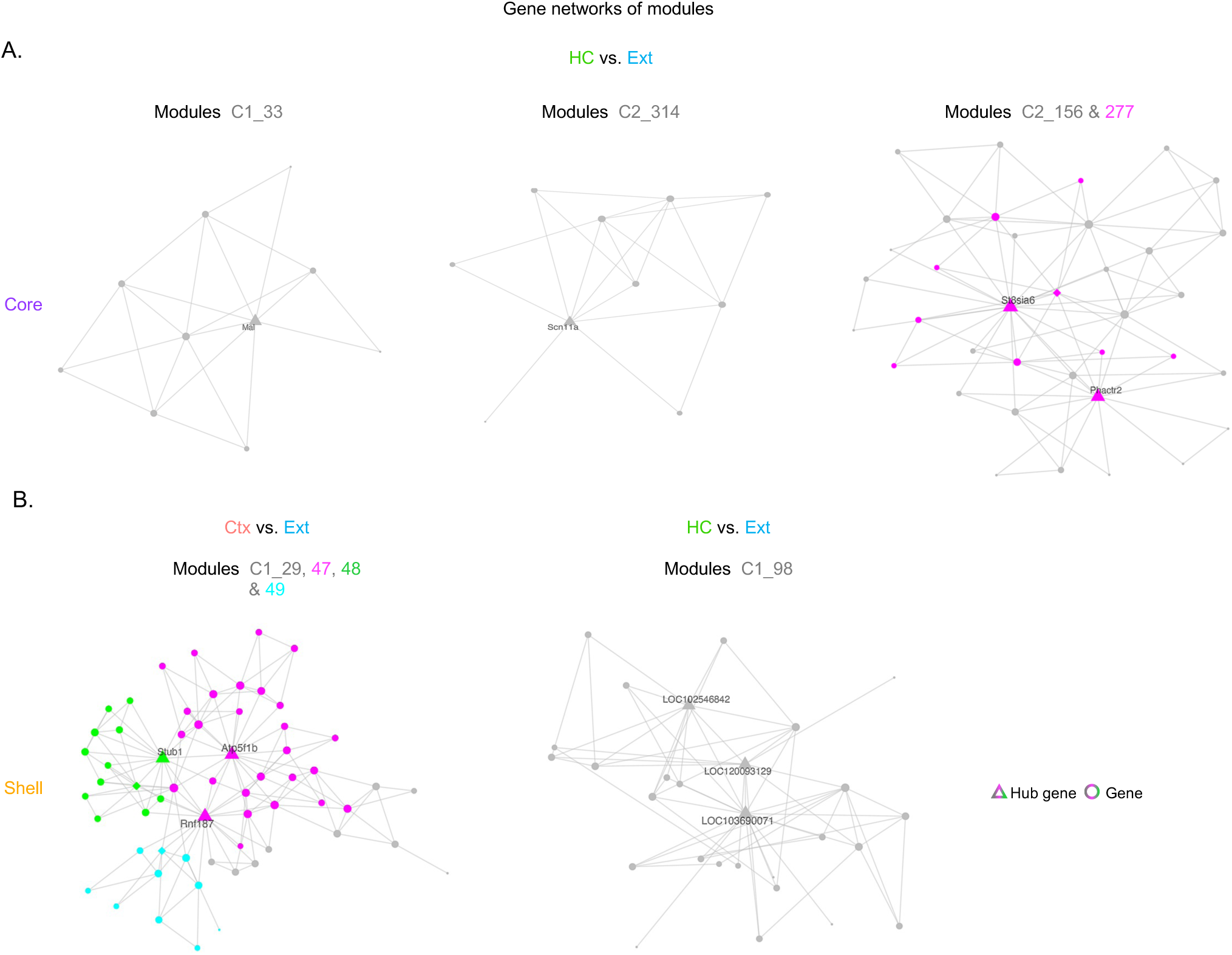
Additional ARACNE networks from overlapping gene co-expression modules across WD/Ext conditions and NAc subregions. **A-B.** ARACNE plots showing gene network and connectivity within additional overlapping modules across WD/Ext conditions and NAc subregions. Hub/driver and connected genes (π = hub gene; λ = gene).

**Table 1 (for Fig. 7).**
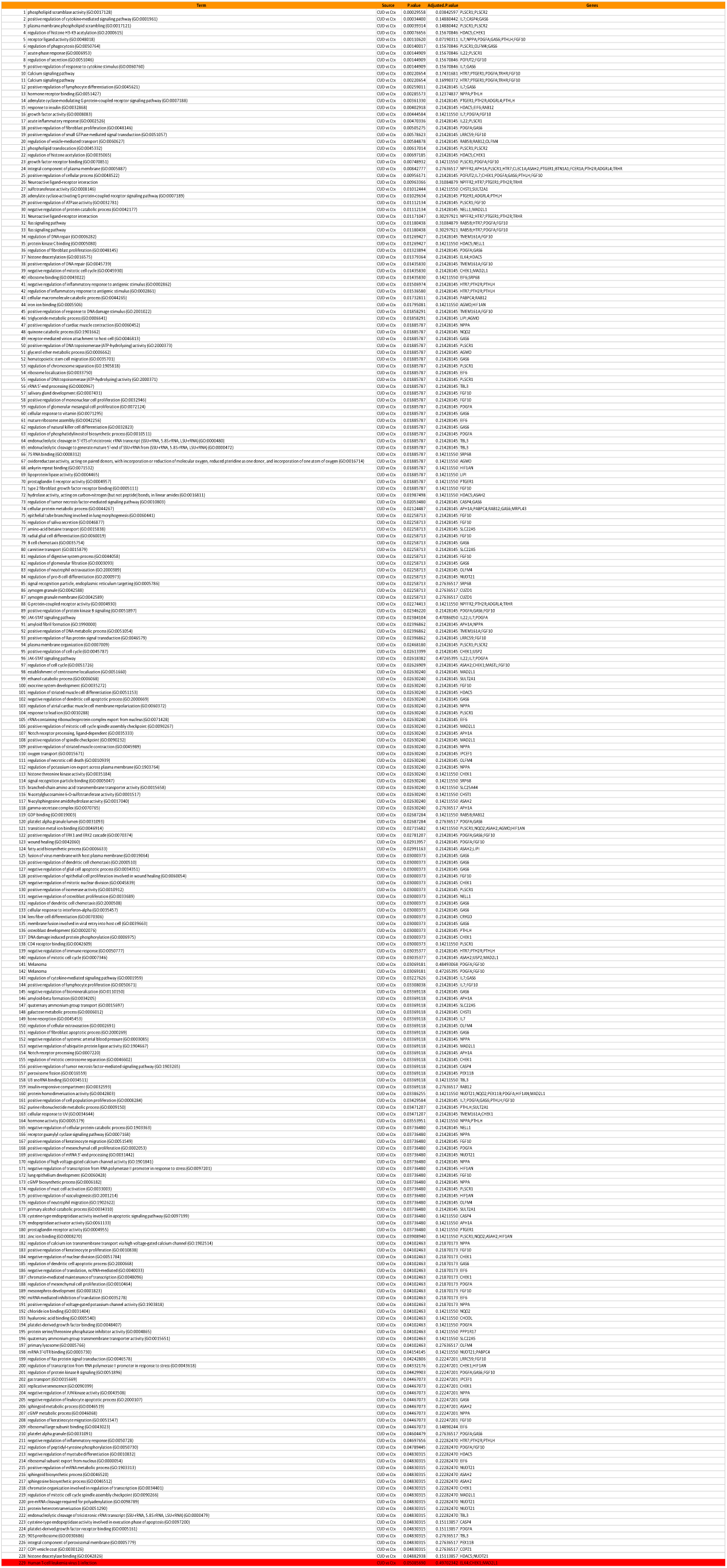
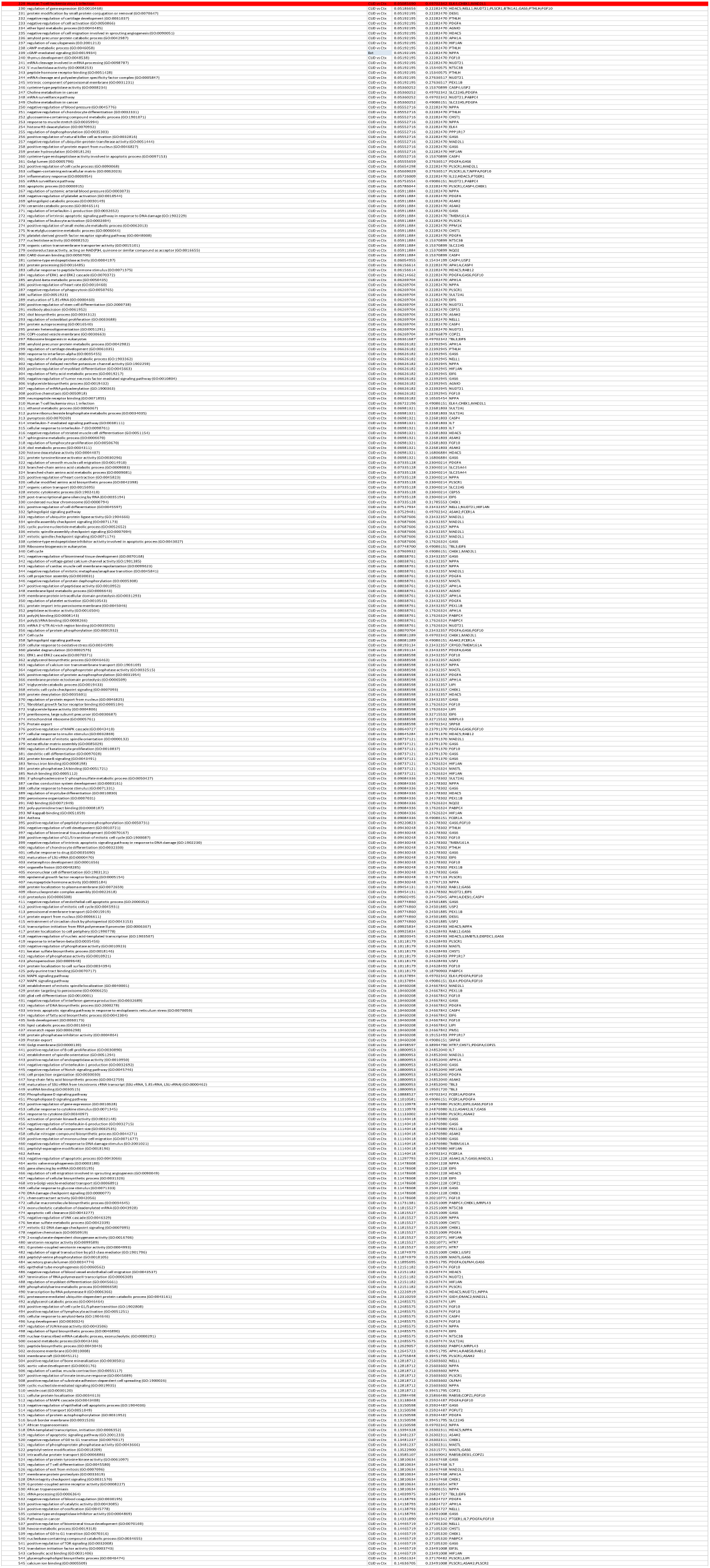
Shared GO terms for transcriptomic regulation in NAc between Ctx withdrawal in rats and human CUD.

